# Persistent glycolysis defines the foreign body response to polymeric implants

**DOI:** 10.1101/2025.02.04.636531

**Authors:** Christian Rempe, Neal Callaghan, Lauren Fong-Hollohan, Sarah Nersesian, Zachary Froom, Kyle Medd, Ibrahim Ahmed, Tobias Karakach, Jeanette E. Boudreau, Michael Bezuhly, Locke Davenport Huyer

## Abstract

Non-degradable polymeric implantable medical devices are a mainstay of modern healthcare but can frequently lead to severe complications. These complications are largely attributable to the foreign body response (**FBR**), which is characterized by excessive inflammation and fibrosis in response to implanted materials. The pathologic mechanisms underpinning the FBR remain elusive; however, metabolism is increasingly regarded as a critical regulator of innate immune function. We conducted comprehensive metabolic profiling of implant-associated macrophages and multinucleated giant cells in response to the subcutaneous implantation of clinically relevant implantable materials in a mouse model of implant fibrosis. Leveraging novel metabolic characterization methods for analysis of both metabolic dependence and enzyme expression in heterogeneous peri-implant tissues, we demonstrate that peri-implant macrophages are glycolytic at least up to six weeks post-implantation. Glycolytically dependent peri-implant macrophages’ expression of glucose transporter 1 (GLUT1) increased temporally and with proximity to the implant-tissue interface. Paired rate-limiting metabolic enzyme expression analysis showed notable increases in biosynthetic pathways (G6PD and ACC1), matched with increased mitochondrial staining intensity in GLUT1^Hi^ cells at chronic timepoints, which were not notable at early timepoints. Notably, we identified a glycolytic dependence of multinucleated macrophages associated with polymeric materials: these cells expressed higher levels of GLUT1 than mononuclear macrophages of comparable metabolic phenotype. Our findings highlight GLUT1-dependent glycolysis as the definitive metabolic system used by peri-implant macrophages and multinucleated cells in the FBR, highlighting this pathway as a potential target for the development of novel therapeutic approaches.

## 1. Introduction

Polymeric implantable medical devices (**PIMDs**) offer mechanical support to replace or reinforce dysfunctional tissue (e.g. breast reconstruction, joint replacements, and surgical meshes). Globally, PIMD use is high, but the clinical impact is compromised by frequent complications due to a unique and chronic inflammatory reaction: the foreign body response (**FBR**)^1^. The FBR underlies implant-associated fibrosis, causing severe pain and impairing function of both the device and host tissues. Moreover, PIMD use has recently been linked with severe systemic illness; textured silicone breast reconstruction has been associated with a 68-fold increase in the risk of anaplastic large cell lymphoma (**ALCL**)^2,3^. This underscores the need to better understand the mechanisms underlying implant-associated inflammation and fibrosis.

The FBR is an innate immune reaction, ostensibly driven by a myeloid response to host proteins adsorbed to the implant surface, including damage associated molecular patterns (**DAMPs**), complement proteins, and endotoxin^4,5^. In the early stages of the FBR, macrophages coordinate and drive pro-phagocytic and pro-angiogenic inflammation, consistent with their role during acute wound-healing^6,7^. However, in the latter stages of FBR progression, the persistence of activating signal (i.e. adsorbed protein) at the implant surface inhibits a complete progression to tissue remodelling and homeostatic functions^8^. Instead, protracted innate activation results in persistent signals that drive fibroblast-mediated extracellular matrix (**ECM**) synthesis (e.g. IL-17, IL-6, TGF-β)^9,10^, ultimately promoting sustained pathological fibrosis. The hallmark of the chronic and pathological phases of this response is the fusion of frustrated macrophages into multinucleated cells (**MNCs**). These cells are believed to form due to the persistence of pro-phagocytotic activating signals, however present phenotyping systems fail to capture the complexity of their role in driving the FBR^1,4,5,11,12^.

Immune function and metabolism are closely linked^13^. Different activation states require specific energetic outputs, and recent evidence suggests that metabolic substrates are involved in cell signalling and transcriptional regulation^14,15^. Canonically, pro-inflammatory cells primarily rely on glycolytic adenosine triphosphate (**ATP**) synthesis, while quiescent and regenerative cells rely on ATP production through efficient but specialized oxidative phosphorylation **(OXPHOS)**^16^. Increased glycolytic reliance has been identified in pathological macrophage-mediated fibrotic illnesses associated with chronic inflammation (e.g. renal and pulmonary fibrosis)^17,18^. Further, environmental influences on cellular metabolism (i.e. hypoxia) have been associated with the development and progression of chronic fibrosis^19^. Mounting evidence demonstrates that the adaptation of central carbon metabolism dictates the functional behaviour of cells in various pathologies: namely through the generation of biosynthetic precursors and reactive oxygen species (**ROS**). For example, in cancer, increased glycolytic and pentose phosphate metabolism support cancer cell proliferation and survival through the synthesis of ribose sugars^20,21^. In acute infection, mitochondrial reverse electron transport supports ROS production to inhibit pathogenic growth^22^, and catabolism of L-arginine into nitric oxide (**NO**) can act to directly inhibit pathogens, as well as promote vasodilation and immune migration to sites of injury^23,24^. Accordingly, immunometabolic interventions have attracted growing therapeutic interest^25,26^. In fact, recent evidence suggests that inhibition of glycolysis and mitochondrial OXPHOS can impact FBR-associated inflammation and fibrotic behaviours^27,28^. The role of macrophage metabolic adaptations underlying the unique chronic inflammation of the FBR, however, remains underexplored, offering an avenue to develop novel, precise therapeutic interventions to address complications.

In this study, we used novel metabolic phenotyping tools to achieve single-cell paired analysis of functional dependence and enzymatic capacity of peri-implant macrophages associated with three clinically relevant materials: polyethylene **(PE**), polydimethylsiloxane (**PDMS** – silicone) and polypropylene (**PP**). We identified persistent glycolytic dependence as the definitive metabolic system in polymer implant-associated macrophages, which localized closely to the implant interface. Temporal adaptations in metabolic enzyme capacity highlighted a unique chronic metabolic phenotype which correlated with the upregulation of signals associated with pathological wound healing in cells where the capacity for glucose import was highest. Each material demonstrated MNC fusion, and these cells were definitively glycolytic in phenotype. We conclude that glycolytic macrophages represent a promising target for therapeutic strategies to mitigate the FBR.

## 2. Results

### 2.1. Fibrotic capsule formation and immunological dynamics are material-independent

To assess material dependence in PIMD associated FBR inflammation and fibrosis, we implanted PE, PP, or PDMS discs subcutaneously in wild-type C57BL/6 mice, and monitored the temporal dynamics of fibrotic capsule formation and associated immune cell behaviour **(Figure 1a)**. Here, implants developed a collagenous fibrotic capsule of comparable thickness, assessed using Masson’s Trichrome staining for each material at 3– and 6-weeks post implantation (**WPI**) **(Figure 1b)**. Capsular thicknesses did not significantly differ between materials at either timepoint **(Supplementary Figure 1)**. Bulk expression of ECM-associated genes (*Col1a1, Col3a1, Fn1*) in capsular tissues supported consistent trends in material-associated fibrosis over time (**Figure 1c**). Expression of rigid scar tissue-associated collagen type I (*Col1a1)*^29^ was upregulated for each material at 6-WPI when compared to 3-WPI (PE: *p* = 0.036, PP: *p* = 0.039), however differences did not reach significance for PDMS-associated tissues (*p* = 0.36). Transcript expression of the pathological fibrosis-associated factor fibronectin-1 (*Fn1)*^30^ was upregulated at 6-WPI when compared to 3-WPI, however differences were only significant for PP-associated tissues (PE: *p* = 0.23, PDMS: *p* = 0.52, PP: *p =* 0.0022). Early scar-associated collagen type III (*Col3a1*)^29,31^ expression was reduced at 6-WPI when compared to 3-WPI in PDMS-(*p* = 0.0089) and PP-(*p =* 0.0099), but not PE-(*p* = 0.19) associated tissues.

**Figure 1:**
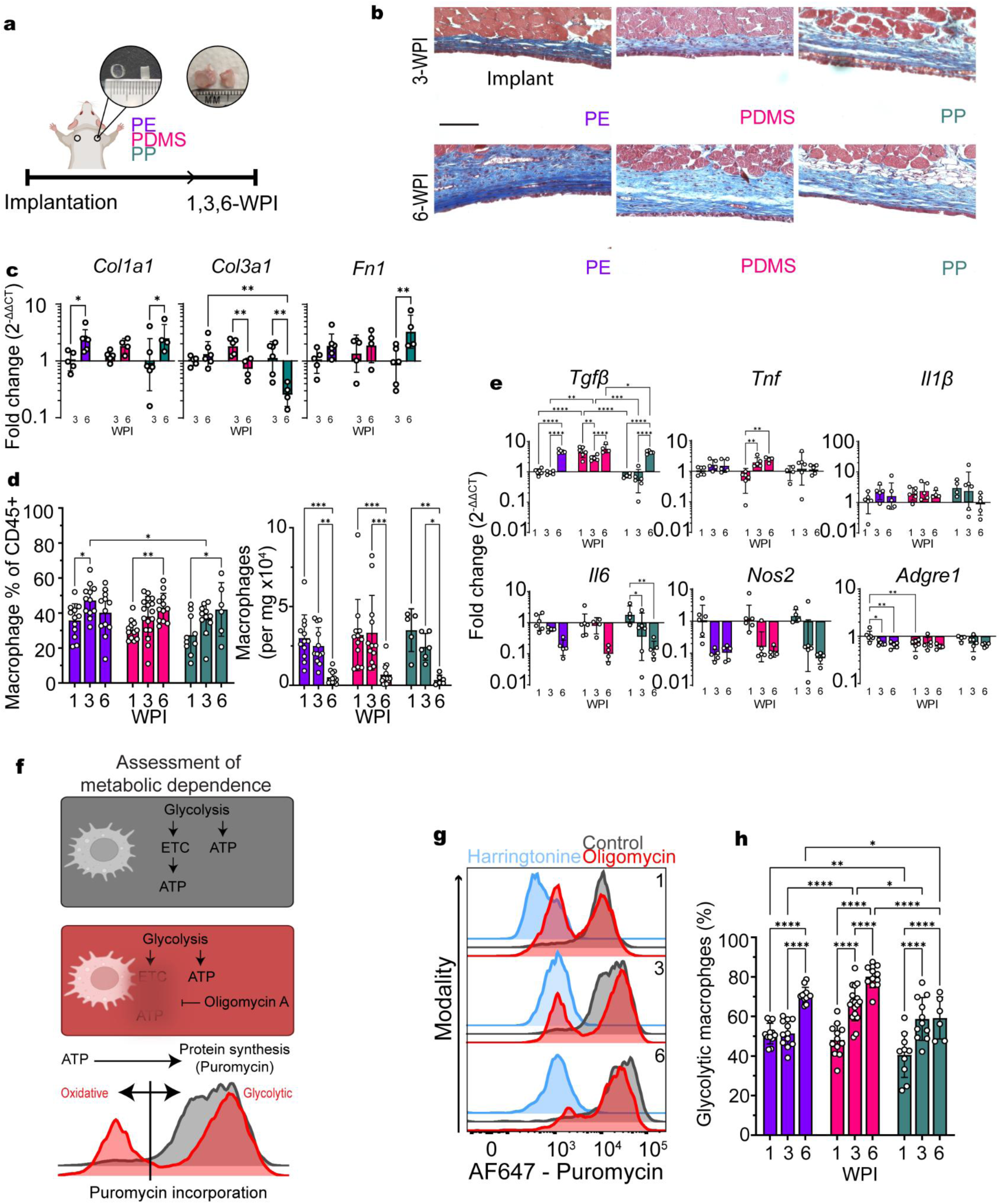
Subcutaneous polymeric implant-associated fibrosis is defined by persistent macrophage dependence on glycolysis. a, Summary of implantation approach: cylindrical polymer discs (PE, PP, or PDMS) were placed subcutaneously (dorsal, subscapular) in male mice for up to 6 weeks. **b-c,** Polymer implants promote development of a fibrotic tissue capsule non-specific to material formulation. **b,** Representative Masson’s Trichrome staining of the implant-associated fibrotic capsule (blue: collagen, red: muscle, black: nuclei) at 3– and 6-WPI. Scale bar = 100 µm. **c,** RT-qPCR of bulk capsular tissues for ECM-associated genes at 3– and 6-WPI. N = 4-6. **d, e** Capsular tissue is macrophage rich with temporal shifts in signaling. **d,** Macrophage frequency of leukocytes and counts per mg of tissue 1-, 3– and 6-WPI. N = 6-17. **e,** Bulk expression of pro-inflammatory and pro-fibrotic signaling factors in capsular tissue 1-, 3– and 6-WPI. n = 4-6. **f-h,** Peri-implant macrophages primarily rely on glycolytic utilization. **f,** Outline of metabolic profiling approach (adapted fromArgüello et al., 2020): alterations to protein synthesis rate (measured using the incorporation of antibody-labeled puromycin to new protein strands) are used as a proxy of impacted ATP production following the addition of specific metabolic inhibitors; oligomycin A targets oxidative phosphorylation while harringtonine targets protein synthesis (negative control). **g,** Representative histograms of temporal puromycin expression in PDMS-associated macrophages. **h**, Temporal frequency of glycolytic (Puromycin^Hi^) macrophages (% of total macrophages) for each material. N = 6-18. Significance determined by two-way ANOVA with Tukey’s multiple comparisons test (**c-h**), and indicated by * *p* < 0.05, ** *p* < 0.01, *** *p* < 0.001, **** *p* < 0.0001.

To determine the temporal immune dynamics of the peri-implant fibrosis, we dissociated capsular tissues into single-cell suspensions and performed multi-parameter flow cytometry. Peri-implant cells were primarily immune myeloid (CD45+, CD11b+) up to 6 WPI (**Supplementary Figure 2a-b**), with no significant difference between material types. Macrophages (CD45+, CD11b+, Ly6G-, F4/80+) represented the largest portion of peri-implant leukocytes and increased in frequency with time (PE: 35.7 ± 9.5%, PDMS: 30.3 ± 5.6%, and PP: 27.3 ± 11.2% at 1-WPI compared with PE: 40.1 ± 12.7%, PDMS: 43.0 ± 8.2% and PP: 42.0 ± 15.3% of cells at 6-WPI, *p* < 0.0001) **(Figure 1d)**. Despite increases in relative macrophage frequencies, cell counts decreased over time (PE: 3003 ± 1465, PDMS: 3188 ± 2264 and PP: 3497 ± 1357 macrophages mg tissue^−1^ at 1-WPI compared with PE: 515 ± 396, PDMS: 677 ± 685 and PP: 341 ± 306 at 6-WPI, *p* < 0.0001) (**Figure 1d**). Neutrophils were also abundant in the peri-implant space, where we observed them to be most frequent at 1-WPI with abundance dropping over time **(Supplementary Figure 2c)**.

Bulk expression of inflammatory and pro-fibrotic transcripts presented similar temporal trends for each material (**Figure 1e**). Transcription of interleukin-6 (*Il6)* and nitric oxide synthase (*Nos2)* (inflammatory mediators associated with fibroblast activation and wound repair)^32–36^ were reduced over implantation duration (*Il6: p* = 0.0002, *Nos2: p =* 0.0052). Transcription of inflammasome-associated pro-interleukin-1β (*Il1β*) was unchanged over time. Transcription of tumor necrosis factor (*Tnf*), an inflammatory mediator associated with fibrotic progression^37^, was similarly unchanged except for observed increases in expression for PDMS-associated tissues from 1-to 6-WPI (*p* = 0.0021). Transcription of transforming growth factor-beta (*Tgfβ1*), a direct activating signal of ECM-producing fibroblasts, increased significantly for each material by 6-WPI when compared with 1-WPI (*p* < 0.0001). The transcription of *Adgre1* (encoding the macrophage marker F4/80) decreased over time for PE-associated tissues (*p* = 0.0044), in line with observed temporal differences in the relative abundance of CD45+ cells (% of live cells) (**Supplemental Figure 2a**).

### 2.1. Glycolysis remains the dominant metabolic phenotype of peri-implant macrophages for at least 6-WPI

To assess the metabolic dependence of peri-implant macrophages, capsular tissues were analyzed using methods adapted from SCENITH^TM^ ^38^. Here, impact to protein synthesis rate (quantified by flow cytometry via incorporation of antibody-labeled puromycin into newly synthesized proteins) is used to identify correlated impacts to whole-cell metabolic rate following targeted pathway inhibition. The integration of metabolic profiling approaches into multi-parameter flow cytometry workflows allowed for analyses in heterogeneous *ex vivo* samples, where cell populations could be identified based on their metabolic dependence and subsequently profiled for phenotypic and functional markers. In this study, we compared peri-implant cells treated with oligomycin-A (ATP synthase inhibitor) to cells from the same tissues treated with a vehicle control (dimethylsulfoxide, **DMSO**) **(Figure 1f-g)**. Where oligomycin-A treatment impacted protein synthesis when compared to the control group, cells were considered oxidative-dependent (reliant on ATP through OXPHOS). Here, temporal analysis of capsular tissues from 1-to 6-WPI indicated that a population of macrophages was unimpacted by oligomycin-A treatment (Puromycin^Hi^) when compared with control-treated samples, suggesting a primarily glycolytic dependence for energy production **(Figure 1g)**. In PE-, PDMS– and PP-associated capsular tissues, the proportion of glycolytic macrophages increased with time (PE: 51.3 ± 5.2%, PDMS: 48.2 ± 8.1%, and PP: 40.5 ± 11.3% at 1-WPI compared to PE: 70.8 ± 3.9, PDMS: 80.1 ± 5.6%, and PP: 59.2 ± 10.6% at 6-WPI, p < 0.0001) **(Figure 1h)**.

Assessments 3-WPI presented comparably high glycolytic dependence in female (**Supplementary Figure 3f**) and male (**Figure 1h**) mice (PDMS: 69.2 ± 4.4% and PP: 59.4 ± 6.1% in females compared to PDMS: 66.6 ± 8.9% and PP: 58.7 ± 10.8% in males; *p* = 0.50 for PDMS and *p =* 0.88 for PP).

A primarily glycolytic dependence was observed for macrophages associated with each material over time. We selected a single material (PDMS) to conduct future focused analyses of metabolic and functional behaviours. To appreciate the association of canonical (*M1* and *M2*) markers in peri-implant macrophages with different metabolic dependences over time, we analyzed common phenotypic markers within glycolytic and oxidative PDMS-associated macrophages through direct multiparameter flow cytometry comparison **(Figure 2a)**. Oxidative macrophages demonstrated upregulated expression of pro-regenerative markers typically associated with *M2* phenotypes^39^ (CD163^+^ and CD206^+^) when compared to glycolytic cells. At 1-WPI, 13.6 ± 6.6% and 28.5 ± 7.5% of oxidative cells were CD163^+^ and CD206^+^, respectively. Consistently, at 6-WPI 10.6 ± 3.5% and 28.5 ± 6.5% were CD163^+^ and CD206^+^, respectively. The relative frequency of CD86+ (*M1*-associated involved in costimulatory T-cell activation) cells was significantly higher among glycolytic cells at each timepoint (36.7 ± 6.5% were CD86+ at 1-WPI and 31.5 ± 9.1% at 6-WPI), although its expression was still notable in the oxidative cells (25.2 ± 4.6% were CD86+ at 1-WPI and 20.2 ± 4.1% at 6-WPI, *p* = 0.0002). The frequency of MHC-II+ (*M1*-associated involved in antigen presentation capacity) cells increased temporally in both glycolytic and oxidative populations (*p* < 0.0001), with significantly higher expression noted in glycolytic cells at 6-WPI (62.8 ± 8.1% of glycolytic cells and 57.3 ± 9.4% of oxidative cells were MHC-II+, *p* = 0.0017). The relative frequency of positive CD163, CD206, CD86 and MHC-II expression within metabolic groups were assessed in PP– and PE-associated peri-implant macrophages at a single timepoint (6-WPI) using the same approach **(Supplementary Figure 4)**.

**Figure 2:**
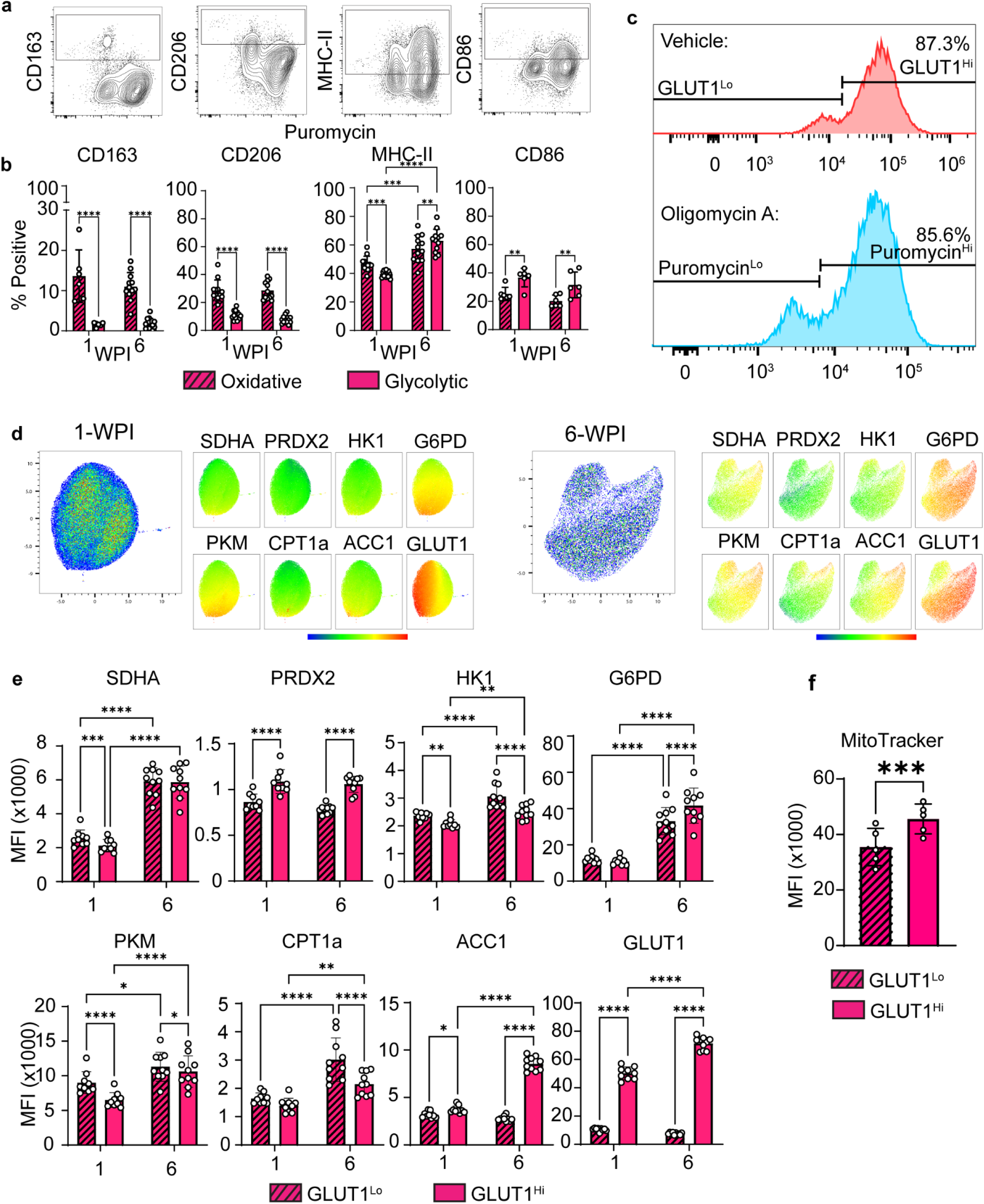
Peri-implant macrophages have distinct and temporally dynamic metabolic profiles a-b, Correlation of canonical macrophage marker expression to glycolytic (Puromycin^Hi^) and oxidative (Puromycin^Lo^) peri-PDMS macrophages. **a,** Representative contour plots (PDMS, 6-WPI) demonstrate gating of canonical markers based on FMO controls. **b,** Frequency of positive expression at 1-WPI and 6-WPI in macrophages with oxidative and glycolytic dependency. n = 6-12. **c,** Expression of GLUT1 directly correlates with glycolytic utilization. Representative histograms of puromycin and GLUT1 expression in matched control– and oligomycin A-treated samples 6-WPI. **d-f,** Expression of key metabolic enzymes in central metabolic processes, which is temporally unique and dependent on central carbon utilization. **d,** Two-dimensional rendering (UMAP) of multi-parameter metabolic enzyme expression in PDMS-associated macrophages at 1– and 6-WPI. N = 9-10. **e,** Quantification of metabolic enzyme expression (MFI) in GLUT1^Hi^ and GLUT1^Lo^ PDMS-associated macrophages 1– and 6-WPI. n = 9-10. **f,** Quantified MitoTracker^TM^ CMXRos MFI in PDMS-associated GLUT1^Hi^ and GLUT1^Lo^ macrophages at 6-WPI. N = 5. Significance determined by repeated measures two-way ANOVA with uncorrected Fisher’s LSD (**b, e**) and paired t-test (**f**), and indicated by \**p* < 0.05, \*\**p* < 0.01, \*\*\**p* < 0.001, \*\*\*\**p* < 0.0001.

### 2.2. Cells with glycolytic and oxidative reliance show distinct enzymatic metabolic signatures which are temporally dynamic

To correlate the functional metabolic dependence in peri-implant macrophages to their capacity, we combined metabolic dependence profiling with intracellular flow cytometric staining of rate limiting enzymes in key metabolic pathways^40^. Using this paired approach, we observed that high expression of glucose transporter 1 (**GLUT1**), the primary transporter for glucose uptake^41^, was strongly correlated with glycolytic reliance (Puromycin^Hi^) on a single-cell level (**Figure 2b; Supplementary Figure 5**). Here, the proportion of GLUT1^Hi^ macrophages in control treatments did not differ statistically from the Puromycin^Hi^ proportion in matched oligomycin-A treatments (1-WPI: *p* = 0.097, 6-WPI: *p =* 0.087). This relationship established GLUT1 as a marker for macrophages reliant on glycolytic processes, eliminating potential impacts of oligomycin A treatment on metabolic enzyme expression^42,43^. Subsequent investigations of metabolic enzyme expression were conducted in vehicle-treated samples based on GLUT1 expression.

The expression profile of rate-limiting enzymes in PDMS-associated peri-implant macrophages indicated temporal differences in metabolic adaptation. Specifically, uniform manifold approximation plots (**UMAPs**) of expression profiles highlighted an evolution of macrophage clusters with increasingly specialized metabolic behaviour between 1-6 WPI (**Figure 2d**). Further gating of macrophages according to GLUT1 expression highlighted distinct and temporally dynamic enzymatic profiles in GLUT1^Hi^ and GLUT1^Lo^ cells (**Figure 2e; Supplementary Figure 6)**. Succinate dehydrogenase A (**SDHA)**, a multifunctional enzyme involved in both the tricarboxylic acid cycle and electron transport chain demonstrated reduced expression in GLUT1^Hi^ cells when compared to GLUT1^Lo^ cells at 1-WPI (1.20 ± 0.05-fold increase in GLUT1^Lo^ cells, *p* = 0.0005), but differences did not reach significance at 6-WPI (*p* = 0.59). Peroxiredoxin 2 (**PRDX2**, antioxidant) was increased in GLUT1^Hi^ cells at both timepoints (1.26 ± 0.04-fold at 1-WPI and 1.35 ± 0.11-fold at 6-WPI, *p* < 0.0001). GLUT1^Lo^ cells were associated with elevated hexokinase 1 (**HK1**) which catalyzes the first reaction in glycolysis under homeostatic conditions (1.13 ± 0.05 and 1.21 ± 0.08-fold increases at 1-WPI and 6-WPI, *p* < 0.0001). Glucose-6-phosphate dehydrogenase (**G6PD**), which catalyzes the first reaction in the NADPH-generating pentose phosphate pathway, was upregulated in GLUT1^Hi^ cells at 6-WPI (1.26 ± 0.14, *p* < 0.0001) but not at 1-WPI (0.90 ± 0.04, *p* = 0.27). Pyruvate kinase M (**PKM**), which catalyzes the penultimate reaction in glycolysis, showed increased expression in GLUT1^Lo^ cells when compared to GLUT1^Hi^ cells at 1-WPI and 6-WPI (1.39 ± 0.03 and 1.06 ± 0.09, *p* < 0.0001). GLUT1^Lo^ cells were associated with upregulations in mitochondrial fatty acid import, mediated by carnitine palmitoyltransferase 1a (**CPT1a**) (1.19 ± 0.09 and 1.36 ± 0.09-fold increases at 1-WPI and 6-WPI, *p* < 0.0001). Acetyl-CoA carboxylase 1 (**ACC1**), which catalyzes the first reaction of fatty acid synthesis, was significantly upregulated (3.13 ± 0.47-fold increase) in GLUT1^Hi^ cells 6-WPI when compared with GLUT1^Lo^ cells 6-WPI (*p* < 0.0001). ACC1 intensity in GLUT1^Hi^ cells was increased at 6-WPI when compared with 1-WPI (1.96 ± 0.16, *p* < 0.0001). Together, these findings support a coordinated intracellular approach in GLUT1^Hi^ macrophages that enables macrophages to adapt and specialize in glycolytic metabolism.

While ATP generation in glycolytic cells is believed to occur independently of mitochondrial OXPHOS, mitochondria are known to drive biosynthetic processes associated with inflammation^44^. To further examine whether observed differences in GLUT1^Hi^ and GLUT1^Lo^ macrophages were consistent with differences in mitochondrial load, MitoTracker^TM^ RedCMXRos staining was performed on capsular tissues, followed by intracellular enzyme staining. Here, glycolytic capacity was correlated with increased mitochondrial staining intensity, where GLUT1^Hi^ PDMS-associated peri-implant macrophages (6-WPI) exhibited significantly higher MitoTracker^TM^ Red CMXRos expression than in GLUT1^Lo^ cells (*p* = 0.0027) (Figure 2f).

### 2.3. Glycolytic macrophages localize at the implant interface

Our results reveal that macrophages with high glycolytic capacity present in response to PIMD. To investigate their spatial organization and potential interactions with the implant material, we utilized multi-parameter immunofluorescent histology (OPAL^TM^) to assess the paired expression of metabolic markers in PE-, PDMS– and PP-associated capsular tissue 3– and 6-WPI. In each material-associated capsule, macrophages (F4/80+) were noted throughout the tissue with focused intensity at the implant interface **(Figure 3a; Supplementary Figure 7)**. We similarly observed a concentrated intensity of GLUT1, lactate dehydrogenase A (**LDHA,** catalyzing the final reaction in committed glycolysis), and cytochrome C (mitochondrial electron shuttle) within the macrophages interfacing directly with each biomaterial, confirming the localization of these metabolically active macrophages at the interface.

**Figure 3:**
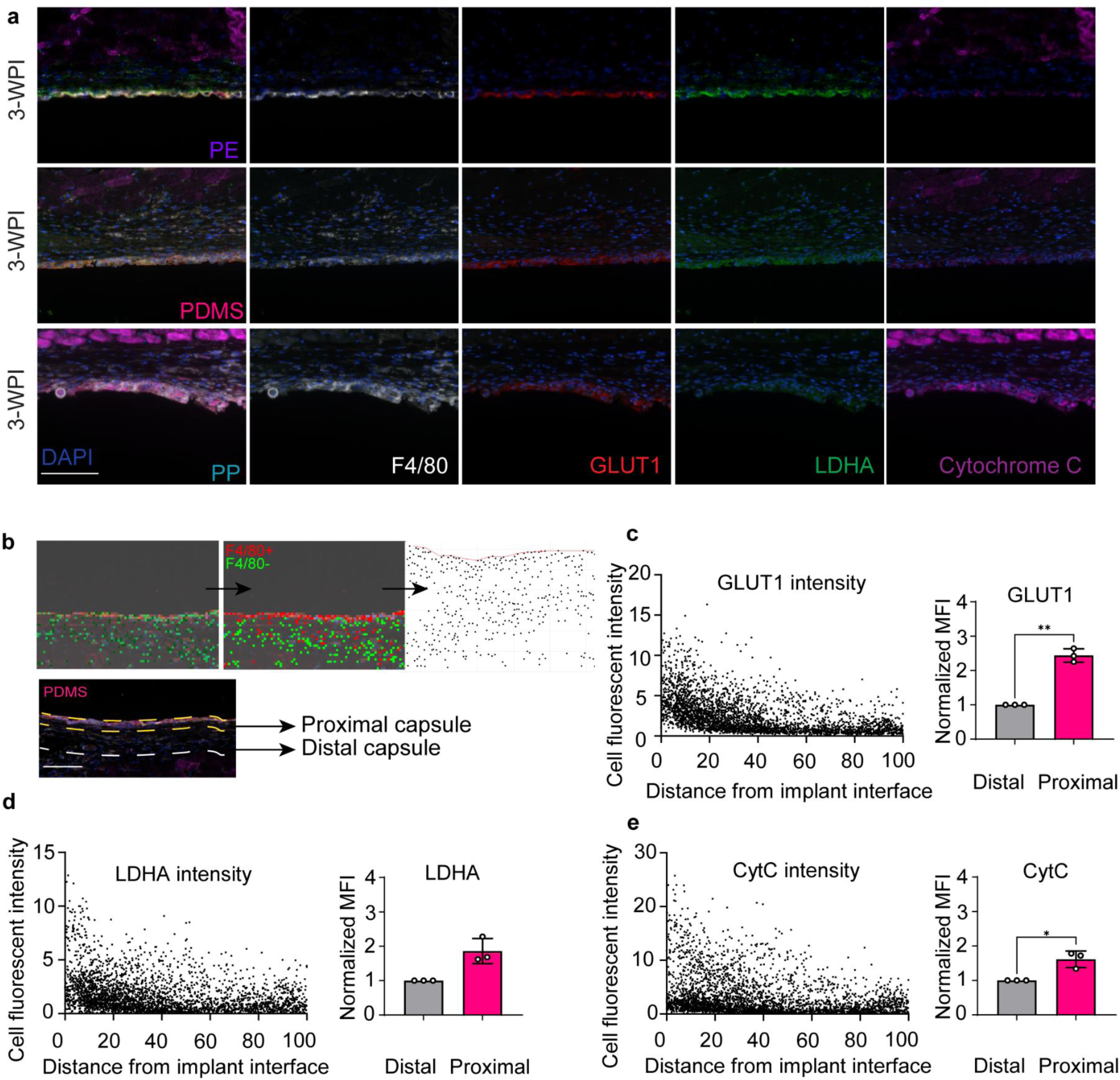
Glycolytic macrophages strongly co-localize with the implant-tissue interface a, Representative Opal^TM^ Immunohistochemical staining of metabolic markers in implant-associated tissues 3-WPI presented as composite images and individual channels for each material. Scale bar = 100 µm. **b – e,** Metabolic enzyme expression (GLUT1, LDHA, and cytochrome C) is increased proximal to the implant. **b,** Automatic spatial metric quantification of metabolic markers in PDMS-associated peri-implant macrophages; representative scatter plots and quantified fluorescent intensity of GLUT1 (**c),** LDHA (**d**) and cytochrome C (**e**) in implant-proximal (<30 µm) macrophages when normalized to their intensity in implant distal macrophages (30-100 µm). N = 3. Significance determined by paired t-test (**c-e**), and indicated by \**p* < 0.05, \*\**p* < 0.01.

We developed a spatial analysis workflow to quantify expression of metabolic markers as a function of distance from the implant surface in PDMS-associated macrophages **(Figure 3b).** Here, cells were individually masked, and their distance from the implant interface and fluorescent intensity of metabolic markers were automatically quantified and recorded (**Figure 3c**). Macrophages were subsequently defined as residing in proximal (<30 μm from the implant interface) and distal (30-100 μm) regions for which the expression of metabolic markers were quantified (**Figure 3c-e**), based on the concentration of macrophages within 30 μm of the interface. Quantified increases in normalized (to distal capsule) median fluorescent intensity (**MFI**) of GLUT1 (2.44 ± 0.20 fold, *p* = 0.0061) (**Figure 3c**), LDHA (1.86 ± 0.36 fold, *p* = 0.055) (**Figure 3d**), and cytochrome C (1.61 ± 0.24, *p* = 0.047) (**Figure 3e**) were observed in the proximal macrophages when compared to the distal macrophages.

### 2.3. GLUT1^Hi^ macrophages co-localize with myofibroblasts

With our observation of GLUT1^Hi^ as a marker of persistent peri-implant macrophages concentrated at the biomaterial interface up to 6-WPI, we further examined their role in the generation of fibrotic capsular tissue. We stained capsular tissues for GLUT1, a marker of myofibroblast activation (α-smooth muscle actin, **a-SMA**), F4/80 and an endothelial cell exclusion marker (CD31) **(Figure 4a)**. Qualitative analysis indicated a-SMA^Hi^ CD31^Lo^ cells (myofibroblasts) were present in the fibrotic capsule, and co-localized with GLUT1^Hi^ macrophages (F4/80+) at both 3– and 6-WPI. This was true for each material at both 3– and 6-WPI, although expression of a-SMA was visibly reduced at 6-WPI **(Supplementary Figure 8)**.

**Figure 4:**
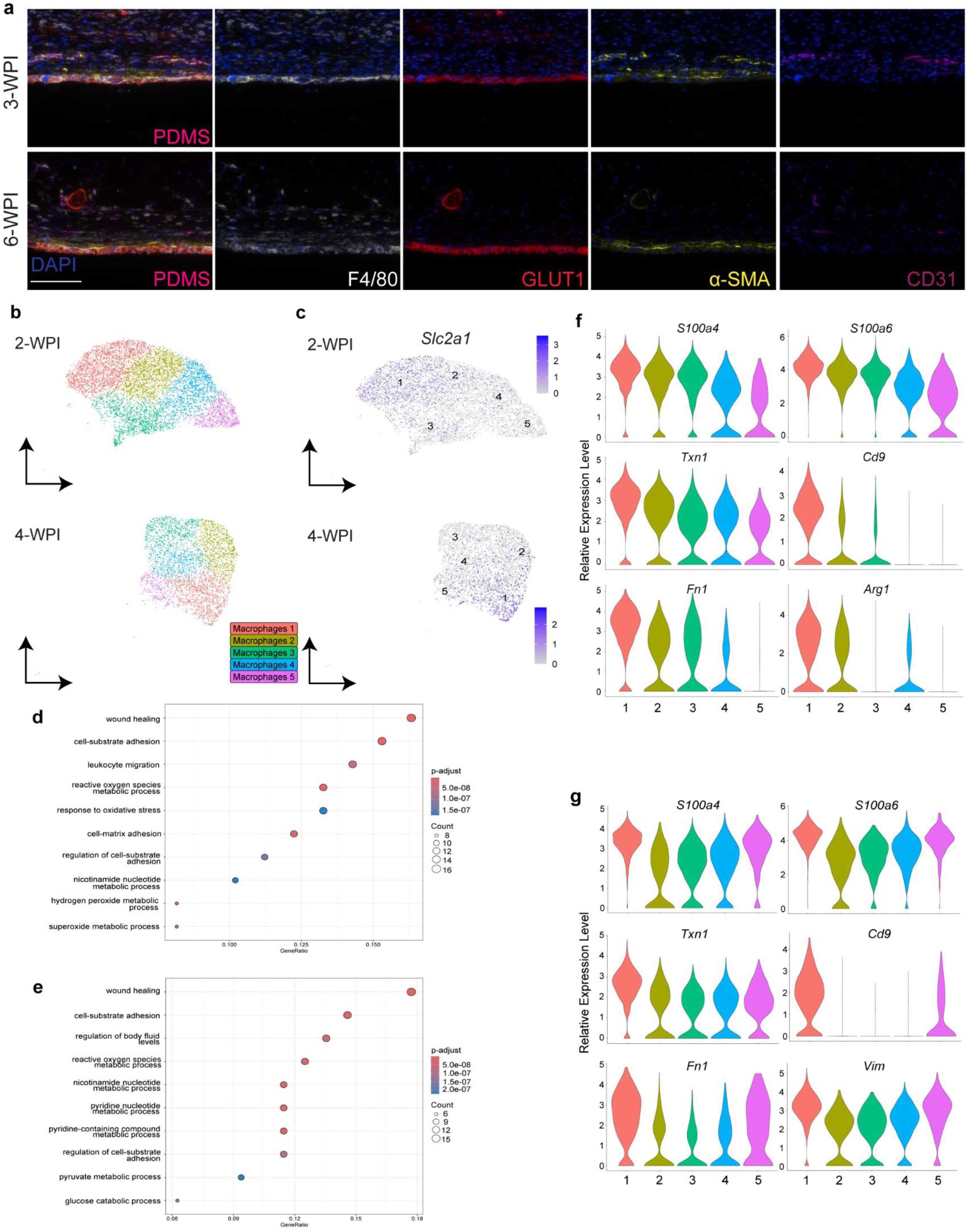
Glycolytic macrophages are associated with pro-inflammatory functions in proximity with activated fibroblasts a, Representative OPAL^TM^ Immunohistochemical staining of fibrotic markers in PDMS-associated tissues 3– and 6-WPI presented as composite images and individual channels. Representative images taken at 40X magnification. Scale bar = 100 µm. **b-e,** Glycolytic macrophages are associated with inflammatory functions. **b,** UMAP of PDMS-associated macrophages from 2– and 4-WPI, manually annotated based on differential gene expression. **c,** Feature map highlighting *Slc2a1* (GLUT1) expression in macrophage clusters at 2– and 4-WPI. Color intensity reflects normalized gene expression levels to the total number of transcripts in each cell. **d-e,** Gene ontology analysis supports wound healing role in macrophages with high glycolytic capacity at 2-(**d**) and 4-WPI (**e**). Top 10 pathways selected based on p-adjusted value and ranked based on gene ratio. **f-g,** Violin plots representing the normalized gene expression levels of six of the top 10 differentially expressed genes in the macrophages clusters 1 at 2-(**f**) and 4-WPI (**g**).

### 2.4. Macrophages with glycolytic transcriptional profiles exhibit functions that drive the pathology of the FBR

To further correlate peri-implant macrophages glycolysis with function, we performed focused analysis of single-cell RNA sequencing datasets in a comparable murine model of subcutaneous PDMS implantation at 2– and 4-WPI^45^ (**Figure 4b**). For each dataset, differentially expressed gene (**DEG**) analysis was performed and five macrophage clusters were manually annotated based on their top 20 DEGs (**Supplemental Figure 9**). Significant upregulation of *Slc2a1* (encoding GLUT1) was observed in macrophage cluster 1 at both 2– and 4-WPI (**Figure 4c; Supplementary Figure 10**). Gene ontology (**GO**) analysis of the top 100 genes in macrophage cluster 1 indicated primary roles in wound healing, mediating oxidative stress and cell adhesion at both timepoints (**Figure 4d-e**). At 2-WPI, remaining macrophage clusters were associated with functions including chemotaxis and migration (clusters 2, 4 and 5) and antigen presentation (clusters 3), whereas at 4-WPI these included responses to inflammatory stimuli (cluster 2), chemotaxis and migration (cluster 3) and antigen presentation (clusters 4 and 5) (**Supplemental Figure 11)**.

Focused analysis of the top 10 DEGs in macrophage cluster 1 at 2– and 4-WPI was conducted to further profile the transcriptional behaviour of highly *Slc2a1*-expressing macrophages associated with silicone implants (**Figure 4f-g**). Here, we observed upregulations in genes associated with diverse pro-inflammatory and pro-fibrotic processes. Cell fusion-associated^46–48^ *Cd9* was the most upregulated DEG at both timepoints. Macrophage activation– and inflammation-associated *S100a6* and *S100a4* were second and third most upregulated at 4-WPI (fourth and 10^th^ at 2-WPI), respectively. The antioxidant *Txn1* (Thioredoxin 1) was the fourth most upregulated DEG at 4-WPI and sixth at 2-WPI. *Fn1* (Fibronectin-1) was the eighth most upregulated DEG at 4-WPI and the second at 2-WPI in the most highly *Slc2a1*-expressing macrophage clusters. Additionally, high *Slc2a1-*expressing macrophages were associated with *Arg1* (Arginase 1) expression at 2– and 4-WPI, representing the 9^th^ and 11^th^ most upregulated DEGs respectively, and with *Vim* (Vimentin) expression at 4-WPI (10^th^ most upregulated DEG).

### 2.5. Multinucleated cells are glycolytic

MNCs are a hallmark of the FBR and well associated with negative clinical outcomes. Histological staining confirmed the presence of MNCs at the implant interface (**Figure 5a**), and qualitative assessment of GLUT1 staining intensity in MNCs indicated strong expression consistent with high glycolytic capacity. Representative overlayed scatter plots (FSC vs. SSC) of oligomycin A-treated peri-implant macrophages gated on Puromycin^Hi^ expression for glycolytic utilization highlighted increases in size within the glycolytic macrophages over time, which was not apparent in the oxidative PDMS-, PE– and PP-associated macrophages **(Figure 5b, Supplemental Figure 12)**. To further assess the role of glycolysis in MNCs, peri-implant tissue samples were analyzed using AMNIS® ImageStream® flow cytometry at 3– and 6-WPI following metabolic inhibition to pair assessment of metabolic dependence with morphological characteristics. ImageStream® analysis of peri-implant macrophages (F4/80+) confirmed an average increase in size (point of maximum thickness) in the glycolytic cells when compared with the oxidative cells at both 3-(size differences for PE: 1.8 ± 0.3µm, PDMS: 2.8 ± 0.5 µm and PP: 2.1 ± 0.4 µm, *p* < 0.0001) and 6-(size differences for PE: 3.5 ± 0.6µm, PDMS: 2.9 ± 0.6 µm and PP: 2.3 ± 0.3 µm, *p* < 0.0001) WPI (**Figure 5c**).

**Figure 5:**
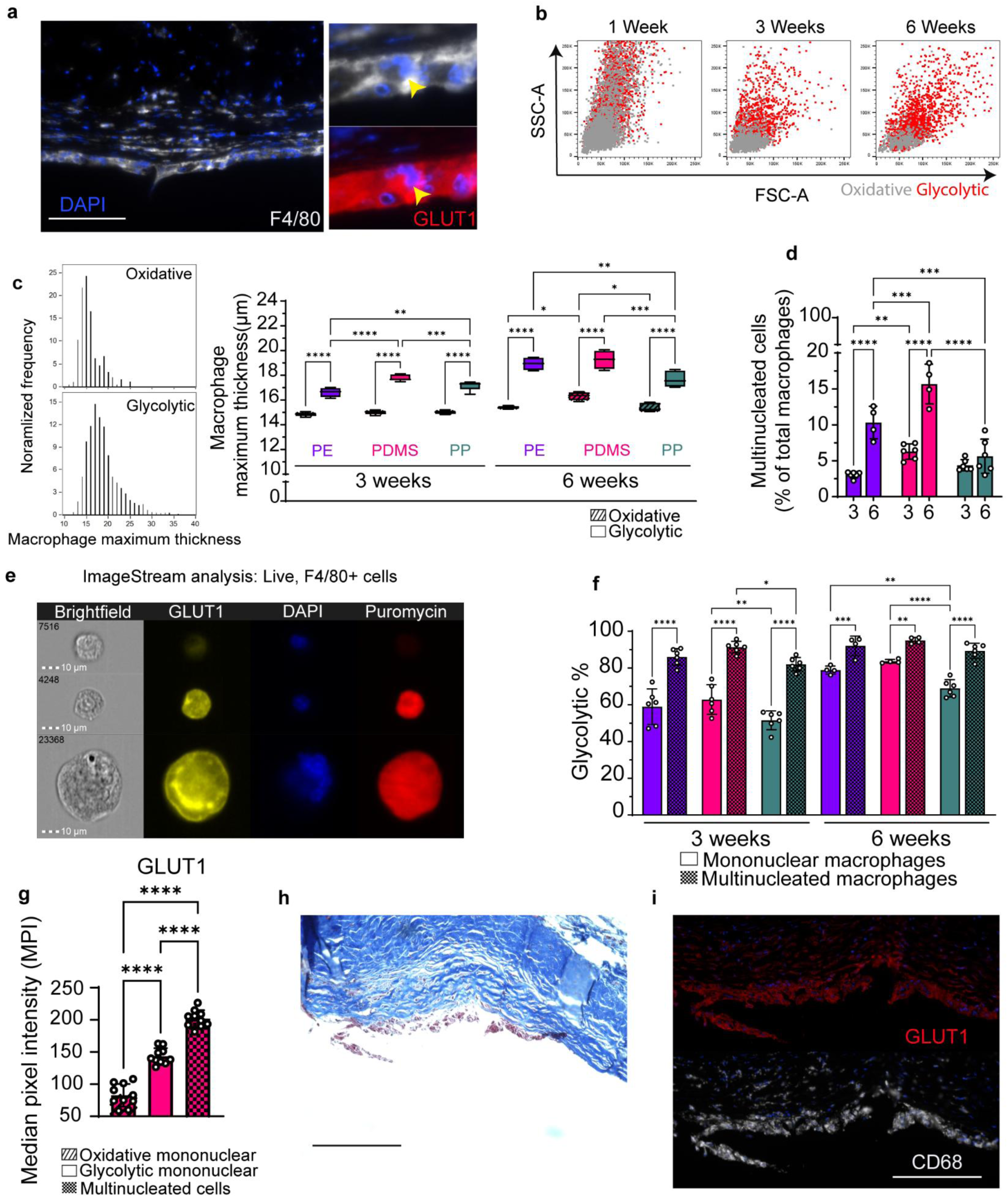
Glycolysis underlies the function of multinucleated peri-implant macrophages. **a**, Representative immunofluorescent staining of PDMS-associated tissues 6-WPI highlights high GLUT1 expression in a multinucleated macrophage at the implant-tissue interface. Scale bars = 100 µm. **b-c,** Glycolytic macrophages are larger than oxidative counterparts and increase in size over time **b,** Representative overlaid scatter plot (FSC-A: size, SSC-A: granularity) of glycolytic (red) and oxidative (gray) PDMS-associated macrophages at 1-, 3– and 6-WPI. **c,** Quantification of glycolytic and oxidative peri-implant macrophages maximal image diameter (Thickness Max) using ImageStream approaches. N = 4-6. **d,** Multinuclear macrophages are present and increase in frequency from 3-to 6-WPI in PE, PDMS, and PP associated capsular tissue. N = 4-6. **e-g,** ImageStream analysis confirms glycolytic utilization and high GLUT1 expression in multinucleated peri-implant macrophages (F4/80^+^). **e,** Representative images from ImageStream® flow cytometry of GLUT1, DAPI (nuclei), and Puromycin expression in mononuclear oxidative, mononuclear glycolytic and multinucleated PDMS-associated macrophages. **f,** Frequency of glycolytic multinucleated and mononuclear peri-implant macrophages. N =4-6. **g,** GLUT1 MPI in multinucleated, mononuclear oxidative and mononuclear glycolytic PDMS-associated macrophages. N = 10. **h-i,** Qualitative assessment of clinical capsular tissues identifies capsular macrophages with high glycolytic capacity. **h,** Masson’s Trichrome and **i,** OPAL^TM^ Immunohistochemical staining (including CD68 and GLUT1) of clinical SBR-associated capsular tissues. Imaged at 20X magnification. N = 1. Scale bar = 100 μm. Significance determined by repeated measures two-way ANOVA with Tukey’s multiple comparisons test **(c, f**), two-way ANOVA with Tukey’s multiple comparisons test (**d**), and repeated measures one-way ANOVA **(f)**, and indicated by *\*p* < 0.05, *\*\*p* < 0.01, *\*\*\*p* < 0.001, *\*\*\*\*p* < 0.0001.

Leveraging image masking for multinuclearity to analyze the frequency and phenotype of MNCs (**Supplementary Figure 13**), we identified increased rates of MNCs (>1 nucleus) over time for each material from 3-to 6-WPI (*p* < 0.0001 for PE and PDMS), however differences did not reach significance for PP-associated tissues (*p* = 0.22) **(Figure 5d)**. PDMS-associated macrophages showed the highest multinucleation (15.7 ± 2.8% of macrophages were multinucleated 6-WPI), followed by PE: 10.3 ± 2.3% and PP: 5.6 ± 2.4% at 6-WPI. Each material exhibited increased rates of multinucleation at 6-WPI when compared to 3-WPI (PE: 3.1 ± 0.4%, PDMS: 6.3 ± 1.0% and PP: 4.8 ± 0.8%, *p* < 0.0001). MNCs at both 3– and 6-WPI were largely unimpacted by oligomycin A treatment and expressed high levels of GLUT1 (**Figure 5e).** Subsequent gating of puromycin incorporation intensity in MNCs indicated high levels of glycolytic dependence at both 3-(PE: 86.0 ± 4.5%, PDMS: 91.4 ± 3.1% and PP: 82.0 ± 3.6%) and 6-WPI (PE: 91.9 ± 5.3%, PDMS: 94.8 ± 1.7% and PP: 89.1 ± 4.2%) (**Figure 5f**). For each material, glycolytic dependence was significantly greater in MNCs when compared to mononuclear cells (increase of PE: 27.0 ± 11.8%, PDMS: 28.5 ± 7.8% and PP: 30.5 ± 5.8% at 3-WPI and PE: 13.2 ± 6.0%, PDMS: 11.4 ± 1.4% and PP: 20.4 ± 6.5% at 6-WPI, *p* < 0.0001). GLUT1 median pixel intensity (**MPI**) was analyzed in mononuclear cells and compared to MNCs for focused analysis in PDMS-associated macrophages at 6-WPI, to appreciate GLUT1 expression with reduced confounding impact of cell size. GLUT1 MPI was significantly increased in MNCs when compared to glycolytic mononuclear cells (1.4-fold increase), which themselves showed 1.7-fold increases when compared with oxidative mononuclear cells (*p* < 0.0001) **(Figure 5g)**.

### 2.6. Clinical silicone implants present with fibrotic capsules associated with glycolytic MNCs

To investigate the metabolism of implant-associated macrophages in clinically relevant samples, we analyzed human silicone breast reconstruction (**SBR**)-associated tissues from a patient undergoing follow-up surgery due to the development of fibrosis. Here, qualitative analysis confirmed the presence of a thick, collagenous peri-implant capsule, with cellularity observed in the implant-proximal regions of the tissues (**Figure 5h**). Strong GLUT1 staining intensity at the tissue-implant interface supported glycolytic metabolism in CD68+ macrophages (**Figure 5i**).

## 3. Discussion

The FBR is an unresolving chronic inflammatory reaction to implanted materials putatively driven by the persistent activation of phagocytes by surface-adsorbed proteins^12^. Chronic immune activity is consistently causative of PIMD complications, including local fibrotic complications and systemic disease (e.g. ALCL)^2,3^. As such, ongoing efforts to characterize this unique inflammatory milieu are required. Previous studies have highlighted the role of macrophages in driving excessive fibrosis during the FBR^1,4,8,9,12^, including their fusion into MNCs that often persist at the implant interface for device lifespan, contributing to pathological outcomes, suggesting that this population may be responsible for the FBR and targets for PIMD tolerance. In this study, we confirmed a consistently macrophage-dominant fibrotic reaction to three clinically relevant polymers and leveraged functional characterization of metabolic profiles to establish novel insight into FBR pathology. Spatial profiling highlighted glycolytic macrophages and MNCs clustering in implant-proximal regions, with transcriptomic analyses supporting active roles for glycolytic cells in mediating pro-fibrotic processes.

Macrophage diversity is achievable through metabolic plasticity, wherein different systems (generally OXPHOS and glycolysis) alter the availability of energy and biosynthetic substrates to achieve function. Pathological alteration of canonical metabolic functions underpins several chronic inflammatory diseases^6,49–52^. Here, we observed a population of macrophages to be metabolically unimpacted by oligomycin A (ATP synthase inhibitor) treatment. These cells were presumed to be glycolysis-dependent, as energetic contributions from amino acid and fatty acid metabolism occur primarily through oxidative phosphorylation and would be impacted by oligomycin A treatment^53,54^. These glycolysis-dependent macrophages persisted into chronic phases of the FBR in a deviation from the progression of canonical wound healing, immediately suggesting a link between pathological metabolic progression and material-associated fibrosis^6,7,55^. These findings were supported by the application of highly novel flow cytometric approaches to assessing the heterogenous FBR immune population in capsular tissue without disrupting their phenotypes with sorting or *in vitro* culturing. The application of functional metabolic profiling^38^ with robust multi-parameter antibody panels allowed us to isolate macrophages and holistically characterize their central carbon metabolic dependence alongside phenotypic features. We directly correlated functional metabolic dependence to specific enzymatic profiles in functionally glycolytic and oxidative cells, identifying GLUT1 expression as a clear indicator of glycolytic macrophages in the FBR. This is consistent with chronically inflammatory macrophage phenotypes elsewhere, including pulmonary fibrosis^51^ and coronary artery disease^52^, and most relevantly with implant fibrosis in a mouse model of muscle regeneration^56^.

Peri-implant glycolytic macrophages exhibited increased GLUT1, ACC1, G6PD, and PRDX2 expression at 6-WPI when compared to 1-WPI, suggesting increased capacity for glucose flux, fatty acid synthesis, pentose phosphate shunting and antioxidant production in chronic phases of the FBR. These differences are consistent with the functional role of glycolytic metabolism in dictating inflammatory functions, supporting the inflammatory and biosynthetic roles of glycolytic macrophages in the FBR, with particularly strong upregulations in the latter stages of the reaction ^21,50,57,58^. Interestingly, glycolytic cells were also associated with elevated mitochondrial expression 6-WPI when compared to oxidative cells, consistent with the previously described reliance on mitochondrial functions for pro-inflammatory ROS production and oxidative stress^59^, a characterized contributing factor to fibrotic development^60^. The strong association of biosynthetic processes (e.g. ACC1^Hi^, G6PD^Hi^, mitochondria^Hi^) and glycolytic metabolism in the FBR at chronic timepoints is noteworthy, and suggests a functional link between cell metabolism and sustained inflammation leading to the development of fibrosis. This observation was supported by the co-localization of a-SMA^Hi^ activated fibroblasts and glycolytic macrophages. Further studies would benefit from examining the contribution of macrophage biosynthetic processes in FBR-associated fibrosis.

The fibrotic reaction to polymeric implants has been previously characterized as having distinct spatial organization^61^. Here, we detected strong localization of glycolytic signatures (GLUT1^Hi^ and LDHA^Hi^) in the implant-proximal regions of the capsule (<30 µm), suggesting the identified glycolytic dependence in implant-associated macrophages is associated with the interface. Interestingly, mitochondrial Cytochrome C was also upregulated with increased proximity to the implant interface, further supporting the increased mitochondrial load in GLUT1^Hi^ cells observed in flow cytometric analyses. Further, we observed macrophage fusion, a hallmark of the FBR, which is now known to be facilitated by hypoxia-associated pathways. Glycolytic stratification with the implant interface suggests a potential role for the hypoxic environment generated by the thick fibrotic capsule underpinning the chronic inflammation of the peri-implant macrophage. Hypoxia has been linked with excessive fibrosis in several pathologies, including bleomycin-induced lung injury, kidney fibrosis, and ischemia^62,63^. However, the direct involvement of hypoxic macrophages as mediators or outcomes of these pathologies, and in the FBR, remains to be interrogated. We demonstrate that these may be related to the development of a glycolytic dependence with associated functional changes.

The functional role of macrophages associated with increased glycolytic capacity was further investigated through analysis of available scRNA-seq datasets in a comparable implant model. Here, GO analysis confirmed that clusters with significantly increased *Slc2a1* expression (GLUT1) were primarily associated with wound healing, mediating oxidative stress, and cell adhesion at both early and late timepoints. This functionality was uniquely distinct from the other macrophage clusters at each timepoint, which supported more canonical inflammatory functions (including chemotaxis and antigen presentation). Interestingly, the expression of *Slc2a1* was relatively low in these clusters, despite their association with inflammatory functions, further emphasizing the importance of glycolytic metabolism in macrophages associated with wound healing in the FBR. *Fn1* was notably upregulated in highly *Slc2a1*-expressing macrophages at both timepoints. With well-accepted pro-fibrotic functions, the expression of *Fn1* highlights a potential direct link between glycolytic metabolism and fibrosis in the FBR^64,65^. Upregulation of *Arg1* was also noteworthy in macrophage cluster 1, representing a top 20 DEG at both 2– and 4-WPI. Arg1 is typically associated with pro-regenerative, oxidative macrophages^39^ but has been correlated with inflammatory activation and glycolytic reprogramming of IL-4 stimulated macrophages^66^. The role of this enzyme in fibrotic progression is further disputed. Some suggest that Arg1^Hi^ macrophages alleviate fibrosis through substrate-stealing^67^. However, other studies indicate that Arg1 mediates the exacerbation of pathology through the production of an ornithine supply critical during collagen synthesis^68^. Interestingly, a recent publication identified high expression of both *Arg1* and *Fn1* as descriptive profibrotic macrophage features, albeit associated with a murine model of myocardial infarction^65^. Together these findings support the role of glycolytic macrophages in driving implant-associated fibrosis.

The functional involvement of fused macrophages in the FBR remains poorly understood due to the incongruence of MNCs with canonical classification systems. We identified glycolysis as the definitive metabolic dependence for implant-associated MNCs, which was accentuated by a higher GLUT1 MPI in MNCs compared to glycolytic mononuclear cells. This supports the notion that glycolytic metabolism is indispensable for MNC function and is consistent with the putative involvement of MNCs in mediating glycolysis-associated functions such as extracellular degradation and inflammation^1^. Metabolic analyses of multinucleated osteoclasts have indicated reliance on glycolysis for bone-resorption functions^69,70^. Interestingly, osteoclast differentiation and fusion are associated with reliance on OXPHOS^69,70^. In our study, 5-15% of MNCs were impacted by oligomycin A treatment, and CD206 was associated with oxidative metabolism. CD206 is the macrophage mannose receptor, and well-documented to be associated with fusion and driving MNC development in the FBR^71^. Taken together, these data could suggest similarly relevant OXPHOS requirements for macrophage fusion, however fusion-associated signals (namely *Cd9*) were observed in macrophages highly expressing *Slc2a1* within the scRNA-seq dataset. Finally, ACC1 expression was noteworthy in glycolytic cells, which were larger than oxidative cells. While ACC1 is generally associated with inflammation (NLRP3 activation via palmitoylation) and is canonically upregulated in glycolytic cells^72^, altered fatty acid metabolism within large cells could indicate shifted membrane dynamics associated with MNC formation and function. Ultimately, further studies would benefit from investigating the metabolic processes associated with MNC fusion and development in the FBR.

This study has identified glycolysis as the definitive metabolic system used by macrophages and MNCs associated with the FBR to PIMDs in mice. Together with our findings identifying GLUT1^Hi^ SBR-associated MNCs and macrophages in clinical samples highlights the potential of metabolically targeted therapeutics to mitigate the FBR and improve clinical outcomes. Although our experimental interventions are reliant on mouse models, which may not effectively recapitulate the complexity and heterogenic severity of the human FBR^45,73^, our preliminary assessment of patient tissues indeed confirms a similar expression of GLUT1^Hi^ macrophages in human SBR-associated capsular fibrotic tissues. Further studies are required to fully determine the role of glycolytic macrophage metabolism in driving the heterogeneity of adverse clinical outcomes in PIMD-associated fibrosis. Further, the role of macrophages in mediating the pathology of the FBR is generally attributed to their orchestration of wound healing and fibroblast communications^74^. Here we have provided evidence of signalling networks associated with glycolytic macrophages, and demonstrated their co-localization with activated fibroblasts. Targeting these interactions could therefore aid in mitigating the pathologic progression associated with PIMD use.

## 4. Methods

### 4.1. Materials

0.5M EDTA (pH 08.0 ± 0.1; Corning 46-034-CI)

1M HEPES (Gibco 15630-080)

1X Antibody Diluent/Block (Akoya Biosciences ARD1001EA)

1X DPBS (Corning 20-031-CV)

1X OPAL^TM^ Polymer HRP MS+RB (Akoya Biosciences ARH1001EA)

1X RPMI 1640 + L-Glutamine (300 mgL^-1^) (Corning 10-040-CV)

1X Plus Manual Amplification Diluent (Akoya Biosciences FP1498)

10% Neutral Buffered Formalin (VWR 16004-128)

Alexa Fluor® 488 Conjugation Kit (Fast) – Lightning-Link ® (ABCAM AB236552)

Alexa Fluor® 700 Conjugation Kit (Fast) – Lightning-Link ® (ABCAM AB269824)

AR6 Antigen Retrieval Buffer (1-5% w/w Sodium Citrate; Akoya Biosciences AR600250ML)

AR9 Antigen Retrieval Buffer (1-2.5% TRIS w/w, <1% EDTA w/w; Akoya Biosciences AR900259ML)

BD Horizon^TM^ Fixable Viability Stain 575V (BD Biosciences 565694)

BSA (Heat shock fraction; Sigma-Aldrich A7906-50G)

Collagenase A (Millipore-Sigma 10103586001)

Chloroform (Fisher Scientific AC423555000)

DAPI (1 mg mL^−1^ stock solution; Thermo Fisher Scientific D1306)

DMSO (Sigma-Aldrich 67-68-5)

DNase I (Millipore-Sigma 10104159001)

DyLight® 405 Conjugation Kit (Fast) – Lightning-Link ® (ABCAM AB201798)

Ethanol (Commercial Alchohols, P016EAAN)

FBS (Heat inactivated, Corning 35-011-CV)

FoxP3 Fixation/Permeabilization Buffer (BioLegend 421403)

FoxP3 Permeabilization Solution (BioLegend 421403)

Harringtonine (ABCAM AB141941)

Histoplast^TM^ PE (Thermo Fisher Scientific 22900700)

Masson’s Trichrome Staining Kit (Polysciences Inc 25088-1)

MitoTracker^TM^ Red CMXRos Dye, for flow cytometry (Thermo Fisher Scientific M46752)

Monocyte Blocker (BioLegend 426102)

Mowiol® 4-88 (Fisher Scientific 9002-89-5)

Oligomycin-A (ABCAM AB143423)

Omni International 1.4 mm Ceramic Bead (Cole-Parmer® 19-645-3)

PE-Cy7® Conjugation Kit – Lightning-Link ® (ABCAM AB102903)

PE-Texas Red Conjugation Kit – Lightning-Link ® (ABCAM AB269899)

Permount® Mounting Medium (Fisher Chemical SP15-500)

Polydimethylsiloxane (PDMS; SYLGARD 184; Dow Chemical GMID 04019862)

Polypropylene (PP; isotactic, avg. MW 220 000; Polysciences Inc. 06536)

Puromycin (Sigma-Aldrich P8833)

Precision Count Beads^TM^ (BioLegend 424902)

Rainbow Calibration Particles, 8 peaks (3.0-3.4 µm, BioLegend 422903)

RNEasy Minikit Plus (Qiagen 74004)

SsoAdvanced Universal SYBR Green Supermix (Bio-Rad 1725274)

SuperScript IV VILO Master Mix (Thermo Fisher Scientific 11756050)

Tris-HCl Buffered Saline + Tween®20 (TBST; 25 mM TRIS-HCl (pH 7.5), 150 mM NaCl, 0.05% Tween®20)

TRIzol^TM^ Reagent (Thiocyanic acid 25-40% w/w, Phenol 30-60% w/w, Ammonium thiocyanate 7-13% w/w; Invitrogen 15596018)

TruStain FcX (BioLegend 101319)

Ultracomp eBeads^TM^ (Thermo Fisher Scientific, 01222242)

Ultra-high MW polyethylene (UHMW-PE; avg. MW 3 000 000 – 6 000 000; Sigma-Aldrich 429015)

Xylene (VWR 6615)

Zombie NIR Fixable Viability Dye (BioLegend 423105)

### 4.2. Implant fabrication

PE and PP were melt-cast using a milled aluminum mold in a 4-ton heat press with upper and lower platens heated at 204° C and 193° C, respectively, to form cylindrical implants (4 mm diameter x 2 mm height). PDMS was generated from Sylgard 184 mixed at a ratio of 10:1 (mass:mass) base:curing agent, degassed under vacuum, and heat-cured at 60°C for 24 hours to form a film of 2 mm height before cutting with 4 mm biopsy punches. After fabrication, implants were chemically sterilized using ethylene oxide prior to implantation.

### 4.3. Subcutaneous implantation

Material implantation (PE, PP, and PDMS implants) was performed in C57BL/6 mice (Charles River Laboratories, male and female, 8-12 weeks of age, approximately 20 g), in accordance with procedures approved by the Dalhousie University Committee on Laboratory Animals (Protocol # 22-027). Anaesthetic was induced and maintained with isoflurane (5% and 2 % respectively, in 1.0 L min^−1^ O_2_). The surgical site was shaved and prepared with Hibiclens® 4% chlorhexidine gluconate, 70% isopropanol, and 10%:1% povidone:iodine. Mice were given subcutaneous analgesia (Meloxicam 5 mg kg^−1^) at the time of surgery. Two incisions were made on the upper back (∼1 cm lateral of the midline), and implants were placed in two subcutaneous pockets caudal to the scapula. Wounds were closed using 9 mm wound clips, which were removed 5 days post-surgery. Mice were euthanized for sampling after 1-, 3-or 6-WPI through anaesthetic overdose and subsequent carbon dioxide inhalation.

### 4.4. Histological sample preparation

Implants, surrounding capsular tissue, and skin were collected, fixed for 48 hours in 30 mL of 10% neutral buffered formalin. Samples were then processed (dehydrated in ethanol and xylene) in a Thermo Fisher Scientific Excelsior AS (A82310100) tissue processor and embedded in paraffin wax. Formalin-fixed, paraffin-embedded (FFPE) samples were prepared using a Leica rotary microtome (RM2255) (4 µm sections), mounted in a warm (34°C) water bath, and dried at 60°C in a VWR 1300U Gravity Convection Oven (11018505) for > 1 hour. Samples were de-paraffinized in xylene and rehydrated stepwise in ethanol before staining.

### 4.5. Fibrotic capsular staining and assessment

Tissue sections were stained using a Polysciences Masson’s Trichrome staining kit according to manufacturer’s instructions. Representative images were captured at 40X (20X for human samples) magnification using a Thermo Fisher Scientific Evos FL Auto 2 microscope. Images for quantification were captured at 10X, and measurements were made in ImageJ2 (U. S. National Institutes of Health, Bethesda, Maryland, USA). Average capsule thickness for each biological replicate was assessed with three measurements (at the most fibrotic, least fibrotic and most representative points) perpendicular to the implant, for each of two-three technical replicates (k = 2-3) per biological sample (total of 9 measurements per biological sample) **(Supplemental Figure 14)**.

### 4.6. RNA extraction and cDNA production

Implants and associated capsular tissue used for RT-qPCR were manually isolated from the cutaneous and fascial tissue layers, flash-frozen in TRIzol (1 mL), and stored at –80°C for subsequent processing. Prior to RNA isolation, tissues were defrosted at room temperature (RT) and disrupted using a VWR Mini Bead Mill (10158-558) in tubes loaded with 1.4 mm ceramic beads for 45 seconds at 5 m s^−1^. Bead-rupting was performed in two bursts (30 s and 15 s) with 30 s of cooling on ice between bursts to reduce sample heating. Genetic material was extracted using 400 mL chloroform and spun at 12,000*g* for 15 minutes (4°C) to achieve phase-separation. RNA was isolated from the aqueous phase and purified using Qiagen RNEasy Minikit Plus according to manufacturers instructions. Sample concentrations were quantified using A260 absorbance obtained with a Thermo Fisher Scientific mDrop plate (N12391) on a Thermo Fisher Scientific VarioSkan Lux absorbance plate reader. Reverse transcription was conducted with 2500 ng of RNA per sample in an Applied Biosystems 2720 ThermoCycler (272S8102236) using ThermoFisher SuperScript IV VILO Master Mix in accordance with the manufacturer protocols.

### 4.7. RT-qPCR

RT-qPCR plates were prepared using 5 ng well^−1^ of cDNA at a reaction volume of 20 μL. Samples were loaded with 10 μL Bio-Rad SsoAdvanced Universal SYBR Green Supermix and primers (forward and reverse) were added to the working solution at a concentration of 0.3 μM. Samples were run using a QuantStudio 3 Real-Time qPCR Instrument (272313147) at an annealing temperature of 60°C and melting temperature of 65°C. Raw Ct values were exported from Design & Analysis v.2.6.0 (Thermo Fisher Scientific, Waltham, Massachusetts, USA). Material-specific relative differences in genetic expression were assessed through a 2^-ΔΔCt^ analysis, where target genes were normalized to the stable housekeeper genes *Rer1* (retention in endoplasmic reticulum sorting receptor 1) and *Canx* (calnexin) in the PE samples from either 3-(**Figure 1.c**) or 1-WPI (**Figure 1.e**). Experiments were performed in duplicate with a threshold of 0.5 CT for technical replicates. A full table of primer sequences used are listed in **Supplemental Table 1**. Primers for genes associated with inflammation and fibrosis were ordered from Thermo Fisher Scientific, sourced from previous publications, and NCBI Primer-BLAST v.2.16.0 (National Library of Medicine, Bethesda, Maryland, USA) was used to verify the reactivity of primers to their respective targets in the *Mus musculus* (taxid: 10090) transcriptome.

### 4.8. Single-cell digestion and metabolic inhibition

To allow for single cell analyses, implants were excised with intact capsules and stored in a 1x Roswell Park Memorial Institute (**RPMI**) 1640 Medium containing L-Glutamine (300 mg L^−1^) and 25 mmol L^−1^ 1x HEPES (**buffered media**). Implant-associated tissues were manually diced into < 2 mm pieces and enzymatically digested (45 min, 37°C) in 8 mL of buffered media containing 0.2 mg mL^−1^ of DNase I and 1.5 mg mL^−1^ of Collagenase A. Digestion was quenched with a matching volume of ice-cold 10% FBS-containing RPMI 1640 media (4°C). Cells were strained and homogenized through a 40 µm PTFE cell strainers prior to use in further analyses.

Samples were treated in accordance with the methods outlined previously for SCENITH^TM^ ^38^. Cells obtained from digestion were placed in a 96-well round-bottom untreated polystyrene plate, split into two groups, and treated with 1 mmol L^−1^ Oligomycin-A or vehicle control (1:1000 DMSO) dissolved in buffered media for 30 minutes. For each experiment, one biological replicate was split three-ways to include a Harringtonine treatment group (2 mg mL^−1^ dissolved in buffered media), which served as a negative control by inhibiting protein synthesis. All samples were subsequently treated with puromycin (10 µg mL^−1^) and cells were incubated for a further 30 minutes at 37° C. Following incubation, cells were centrifuged and washed with 1X Dulbecco’s phosphate buffered saline (**DPBS**) prior to flow cytometric staining.

### 4.9. Flow cytometry staining

Cells were stained with reconstituted Zombie NIR Fixable Viability Dye (1:1000 dilution in 1X DPBS, 30 min, 4°C) or BD Horizon^TM^ Fixable Viability Stain 575V (1:2000 dilution in 1X DPBS, 30 min, 4°C) and washed twice with 1X DPBS solution containing 1% (w/v) BSA and 1 mmol L^−1^ EDTA (**FACS Buffer**). Surface staining was performed with a solution of FACS Buffer, TruStain FcX, and Monocyte Blocker (1:50 dilution each) for 45 minutes, covered over ice.

For MitoTracker^TM^ experiments, cells were then washed twice with 1X DPBS and stained with MitoTracker^TM^ Red CMXRos (1:1000 in 1X DPBS, 30 min, RT).

Following surface staining, cells were washed twice with FACS Buffer, fixed and permeabilized using the BioLegend FOXP3 fixation/permeabilization kit in accordance with manufacturer instructions. Intracellular staining was performed by diluting antibodies with FOXP3 permeabilization buffer containing TruStain FcX (1:50) and Monocyte Blocker (1:50 each). Cells were stained for 30 minutes at RT.

For ImageStream® experiments, cells were washed with FOXP3 permeabilization buffer, following intracellular staining and incubated at RT in DAPI (0.5 mg mL^-1^ dilution in FOXP3 permeabilization buffer) for a further 10 min.

For each experiment, single-stain controls (SSCs) and fluorescence minus-one controls (FMOs) were prepared for each marker. A summary of the antibodies, supplier information and relevant concentrations used are listed in **Supplemental Table 2**. Unconjugated antibodies (anti-G6PD, anti-ACC1, anti-PKM, anti-CPT1a, anti-SDHA) were prepared using Lightning-Link® from ABCAM in accordance with manufacturer instructions. Cell counts were obtained using 50 mL of Precision Count Beads in control-treated or harringtonine-treated samples.

### 4.10. Flow cytometry analyses

Flow samples were read on a BD FACSCelesta^TM^ Cell Analyzer, a BD FACSymphony^TM^ A5 Cell Analyzer, and a Cytek® AMNIS® ImageStream® Mk II Imaging Flow Cytometer (imaging at 40X magnification). Standard flow cytometry analyses were conducted in FlowJo v.10.9.1 (BD Biosciences, Franklin Lakes, New Jersey, USA). To identify macrophages, samples were gated sequentially to exclude post-digestion debris, cell doublets, dead cells, CD45-, CD11b-, Ly6G+ and F4/80-cells **(Supplemental Figure 15, Supplemental Figure 16)**. For each experiment, FMO controls were used to ensure accurate identification of positive staining. UMAPs were generated using default settings (nearest neighbours = 15, minimum distance = 0.5) to conserve balanced global and local structure in the UMAP v.4.1.1 plugin (BD Biosciences, Franklin Lakes, New Jersey, USA) for FlowJo v.10.9.1. For each experiment, replicates were downsampled using DownSample v.3.3.1 (BD Biosciences, Franklin Lakes, New Jersey, USA) and concatenated to ensure equal representation for each sample. Imaging flow cytometry analysis was conducted using IDEAS v.6.2 (Cytek AMNIS, Seattle, Washington, USA). Macrophages were identified by sequentially gating to remove post-digestion debris/cell doublets, dead cells and DAPI-F4/80-cells (**Supplemental Figure 13**). Macrophage size was assessed using the Thickness Max feature, which measures the width of an imaged object mask at its maximum point. Multinucleation was assessed using Threshold (coefficient 86) and Watershed (intensity weighted, line thickness 1, smoothing 4) masking on DAPI and filtered downstream to eliminate erroneously masked single cells (based on Brightfield Minor Axis and Minor Axis Intensity features to distinguish small cells (>20 each)) and clumps (based on Brightfield Shape Ratio (>0.8) and Thickness Max (>20) features to identify large cells with near equivalent width and height) (**Supplemental Figure 13**). Downstream filtering criteria were identified using the feature finder wizard, trained on user-selected MNCs and erroneously masked single-cells and clumps. The quality of multinucleated cells was manually confirmed following upstream filtering.

### 4.11. Opal^TM^ multiplexed fluorescent immunohistochemical staining

Opal^TM^ multiplex fluorescent IHC staining was performed in accordance with manufacturer’s protocols. Briefly, following deparaffinization, slides were fixed in 10% neutral-buffered formalin for an additional 20 minutes, followed by heat-mediated antigen retrieval in a microwave using AR6 or AR9 buffers. Slides were washed (three two-minute washes) with TBST in between each step. Slides were blocked with Akoya Biosciences 1X Antibody Diluent/Block. Primary antibody dilutions were made using the 1X Antibody Diluent/Block. Secondary antibody incubation was performed using 1X Opal^TM^ Polymer HRP MS+RB from Akoya Biosciences. OPAL^TM^ fluorophore dilutions (1:100) were made using Akoya Biosciences 1X Plus Manual Amplification Diluent. A 15 mM DAPI solution was used as a nuclear counterstain. A summary of the antibodies and relevant concentrations used are listed in **Supplemental Table 3**. Slides were mounted using Mowiol® 4-88 mounting medium, stored at 4°C and imaged within 7 days of staining.

### 4.12. Fluorescent immunohistochemical analysis

All Opal^TM^ imaging was conducted using a Mantra^TM^ Quantitative Pathology Imaging Station, using MantraSnap v.1.0.4 (Akoya Biosciences, Marlborough, Massachusetts, USA). Representative images were captured at 40X magnification (20X for clinical samples). Images for quantification were captured at 20X, then processed in inForm v.2.4.8 imaging analysis software (Akoya Biosciences, Marlborough, Massachusetts, USA). Primary-delete controls were included for each marker and used to subtract background fluorescence from the stained samples. Cells were segmented using object-based nuclear masking (DAPI, minimum size of 50 pixels, maximum size of 400 pixels, intensity threshold of 0.17), and F4/80+ cells were identified with user-training **(Figure 3.b)**. Cell coordinates, phenotypes and the fluorescent intensity of markers were exported to RStudio v. 4.3.3 (Posit PBC, Boston, Massachusetts, USA). Dplyr v.1.1.4 (Hadley Wickham & Romain François, 2023), ggplot2 v.3.5.1 (Hadley Wickham, 2016) and reshape2 v.1.4.4 (Hadley Wickham, 2020) packages were used for analyses. (Tissue slides were digitally reconstructed using cell coordinates, and the implant interface was traced with a curve based on the mean X coordinates and maximum (or minimum, depending on the image orientation) Y coordinates for every 30 cells. F4/80+ filtering was conducted using the phenotypic classifications from inForm (F4/80+/-). Euclidean distance of each F4/80+ cell from the interface curve was calculated. Analyses were run in triplicate for each biological replicate (k = 3).

### 4.13. Single-cell RNA-sequencing dataset analyses

Single-cell RNA-sequencing data was retrieved from the publicly accessible Gene Expression Omnibus (GEO) database. Datasets examining subcutaneous silicone implants in the mouse dorsum at 2– and 4-WPI^45^ were used for subsequent analyses. Seurat v.5.1.0 (Butler et al., 2024) package for RStudio (v.4.3.3) was used for downstream scRNA-seq data analyses. Doublets and cells with fewer than 500 unique transcripts, >10% mitochondrial counts and >30-35% ribosomal counts were excluded. After filtering, a total of 10,446 cells at 2-WPI and 6,828 cells at 4-WPI were retained for further analysis. The gene expression for each cell was normalized to the total expression using a global-scaling log-normalization method with a scale factor of 10,000. The first 15 principal components at 2– and 4-WPI were used to cluster cells and generate UMAPs. DEGs with positive expression were determined for each cluster using Seurat’s FindMarkers function with a log2-fold change threshold of >1. Cell clusters were manually annotated based on the top 20 DEGs. A subset consisting of only the macrophage clusters was generated, which included 6297 cells at 2-WPI and 3470 cells at 4-WPI. Differential gene expression (top 20 genes per cluster) analysis was performed on the macrophage subsets using the FindMarkers function. Macrophage clusters in which *Slc2a1* was significantly differentially expressed were identified with a p-value threshold of <0.05. Gene ontology analysis was conducted on the top 100 upregulated DEGs using the clusterProfiler R package v.4.12.6 (Yu et al., 2024). Genes were annotated using the org.Mm.eg.db v.3.19.1 (Carlson, 2024). The top 10 biological processes were determined based on adjusted *p* values and ranked based on gene ratio.

### 4.14. Human sample ethics declaration

Deidentified FFPE SBR-associated capsular tissues were collected from consenting patients receiving SBR revision or removal surgeries in accordance with the Nova Scotia Health Research Ethics Board (**NSHREB**) approved protocols (REB # 1030720).

### 4.15. Data representation and statistical analyses

Graphs were generated and statistical analyses conducted in GraphPad Prism 10 v.4.3.0 (GraphPad Software Inc., San Diego, California, USA). Statistical analyses utilized are noted in the respective figure captions. Figures were assembled in Adobe® Illustrator v.28.5 (Adobe Inc., San Jose, California, USA).

## 5. Author Contributions

CR, NC, and LDH conceived the idea, designed the experiments and analyzed the results. CR, NC, LFH, SN, ZF, KM, IA, and LDH performed experiments and analyzed the results. CR and NC performed animal surgeries. LFH and IA performed single-cell transcriptomic analyses. CR and SN developed and performed spatial analyses. ZF and KM performed RT-qPCR experimentation and analyses. CR wrote the manuscript. TK supervised single-cell transcriptomics, JB supervised spatial analysis, MB supervised fibrotic assessment and clinical studies. LDH supervised the entire project. All authors read the manuscript, commented on it and approved its content.

## 6. Acknowledgements

This research is supported by the Natural Sciences and Engineering Research Council of Canada (NSERC) Discovery Grant (RGPIN-2022-03666), Dalhousie Medical Resarch Foundation – Faculty of Dentistry Early Career Researcher Award, New Frontiers in Research Fund – Exploration Fund (NFRFE-2022-00313), and Canadian Institutes of Health Research Project Grant (PJT – 191692). We acknowledge infrastructure support from the Canadian Foundation for Innovation (JELF 42294) and Research Nova Scotia Research Opportunities Fund (2022-2400). The authors would like to acknowledge the assistance of Riley Arseneau, Jorge Pinzon Mejia, Derek Rowter, Renee Raudonis, Edwin Leong, Natasha Osbourne, Brenden Wheeler, Angela Xu, Stephanie Kansiime, and Julia Harrison.

CR was supported by a CIHR Canada Graduate Scholarship – Masters. ZF was supported by an NSERC Canada Graduate Scholarship – Masters, Nova Scotia Graduate Scholarship, and Killam Predoctoral Scholarship (Masters).

