## Supplemental figures and tables for "Persistent glycolysis defines the foreign body response to polymeric implants"

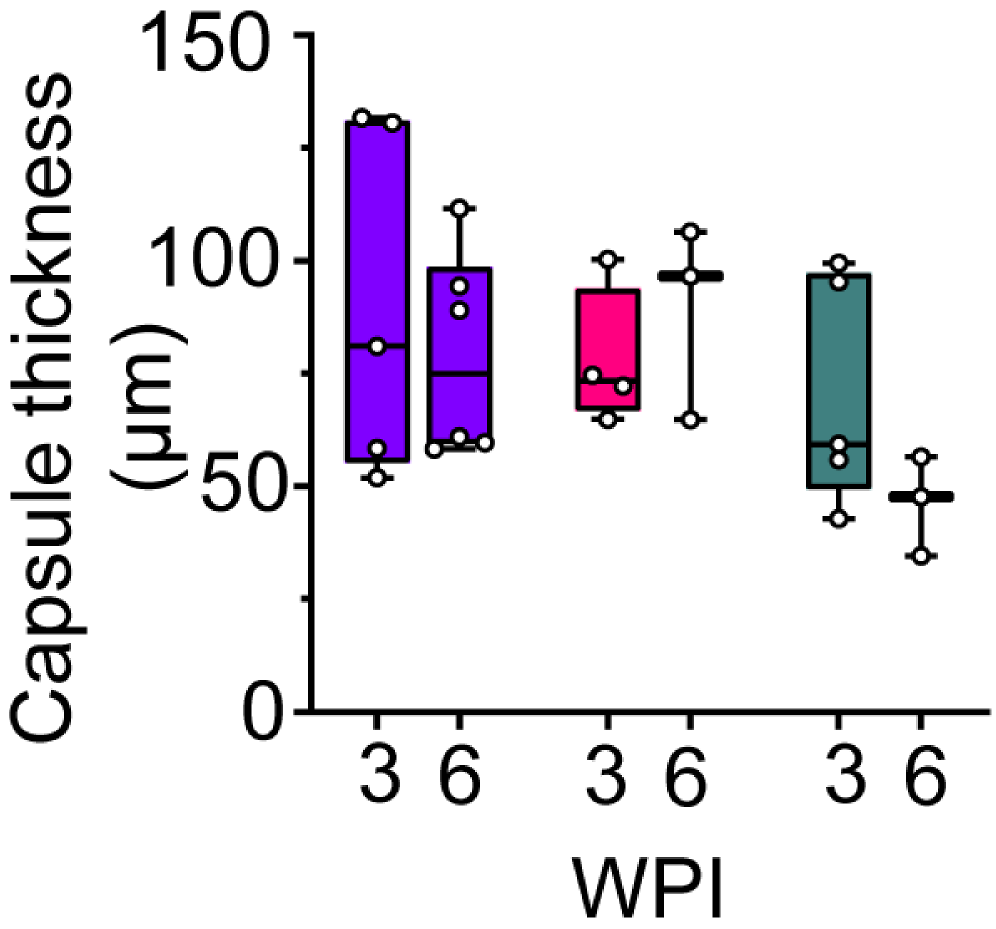


**Supplemental Figure 1:** Fibrotic capsule thickness associated with subcutaneous polymer implantation at 3- and 6-WPI. k = 2-3, n= 3-6. Two-way ANOVA with Fischer's LSD. $*p < 0.05$.

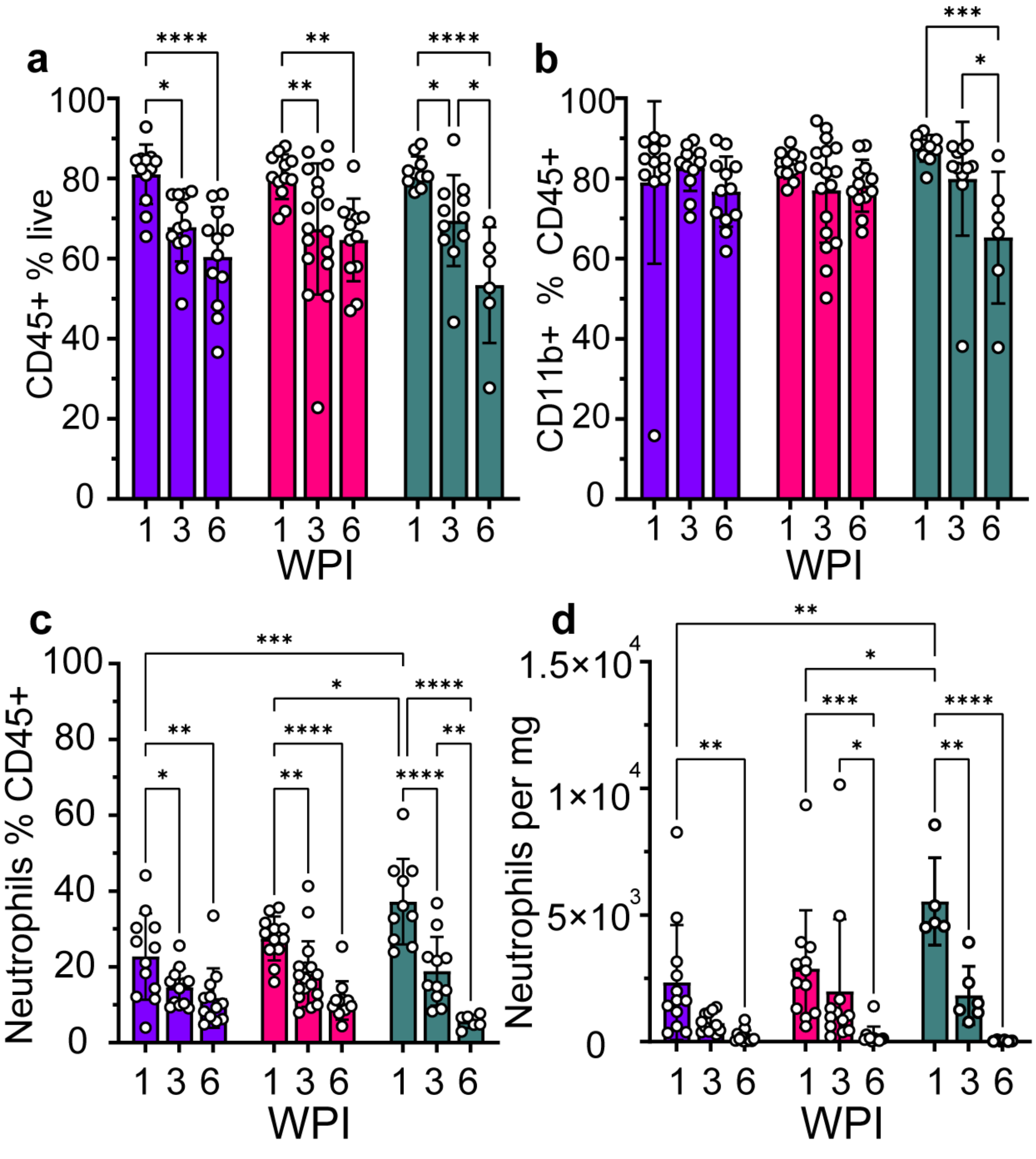


**Supplemental Figure 2: Temporal leukocyte dynamics of the PIMD associated FBR.** Frequency of **a,** total leukocytes (CD45+) **b,** myeloid cells (CD11b+)**,** and **c,** neutrophils (Ly6G+ $Ly6C^{Mid}$). **d.** Neutrophil counts per mg of explanted tissue. n = 6-17. Compared using a regular two-way ANOVA with Tukey's multiple comparisons test. *$p < 0.05$, **$p < 0.01$, ***$p < 0.001$, ****$p < 0.0001$.

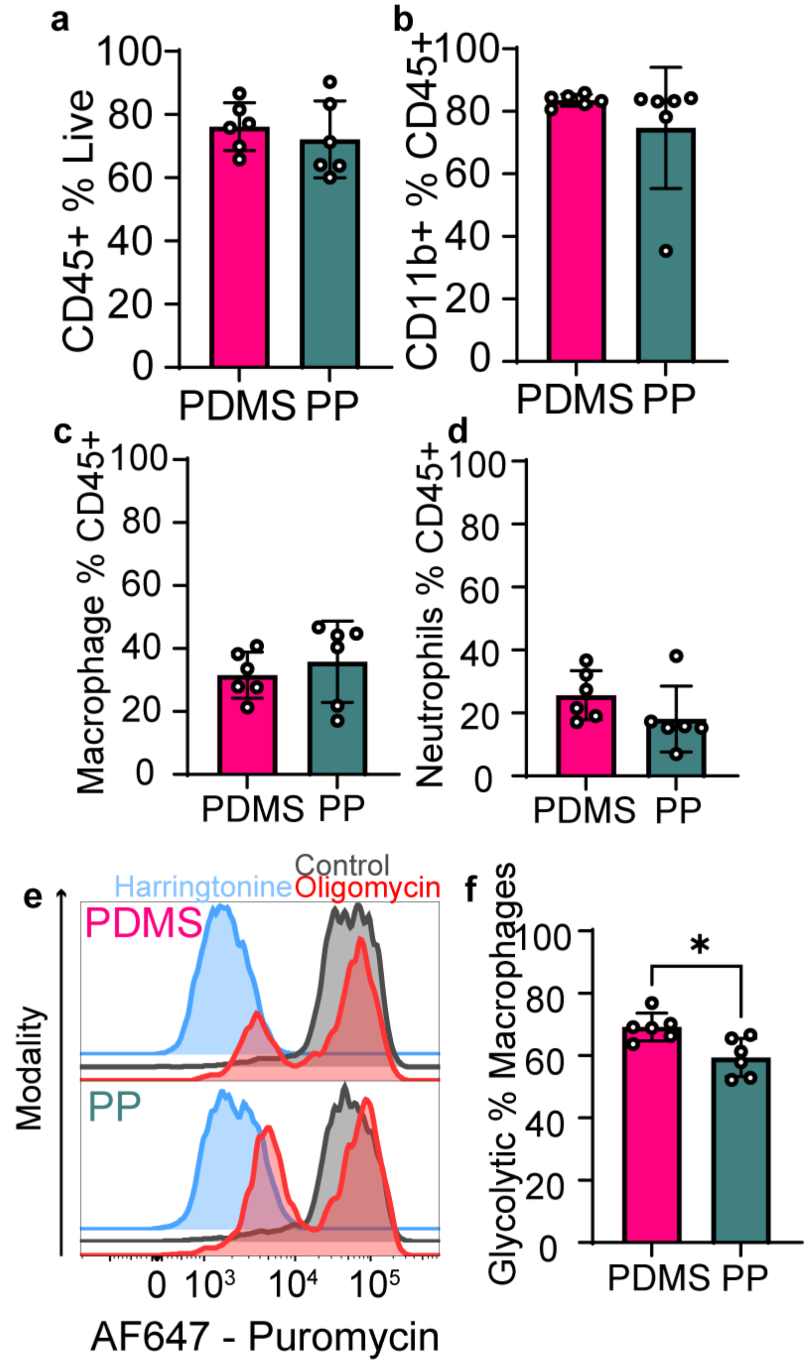


**Supplemental Figure 3: Leukocyte dynamics and central carbon peri-implant macrophage metabolic utilization associated with PDMS and PP in female mice at 3-WPI.** Frequency of **a,** total leukocytes (CD45+) **b,** myeloid cells (CD11b+)**, c,** macrophages (F480+), and **d,** neutrophils (Ly6G+ Ly6C$^{mid}$). **e,** Representative histograms of puromycin expression in peri-implant macrophages associated with PDMS and PP implants. **f,** Frequency of macrophages that are glycolytic (Puromycin$^{Hi}$**)**. n = 6. Compared using an unpaired t-test. *$p$ < 0.05.

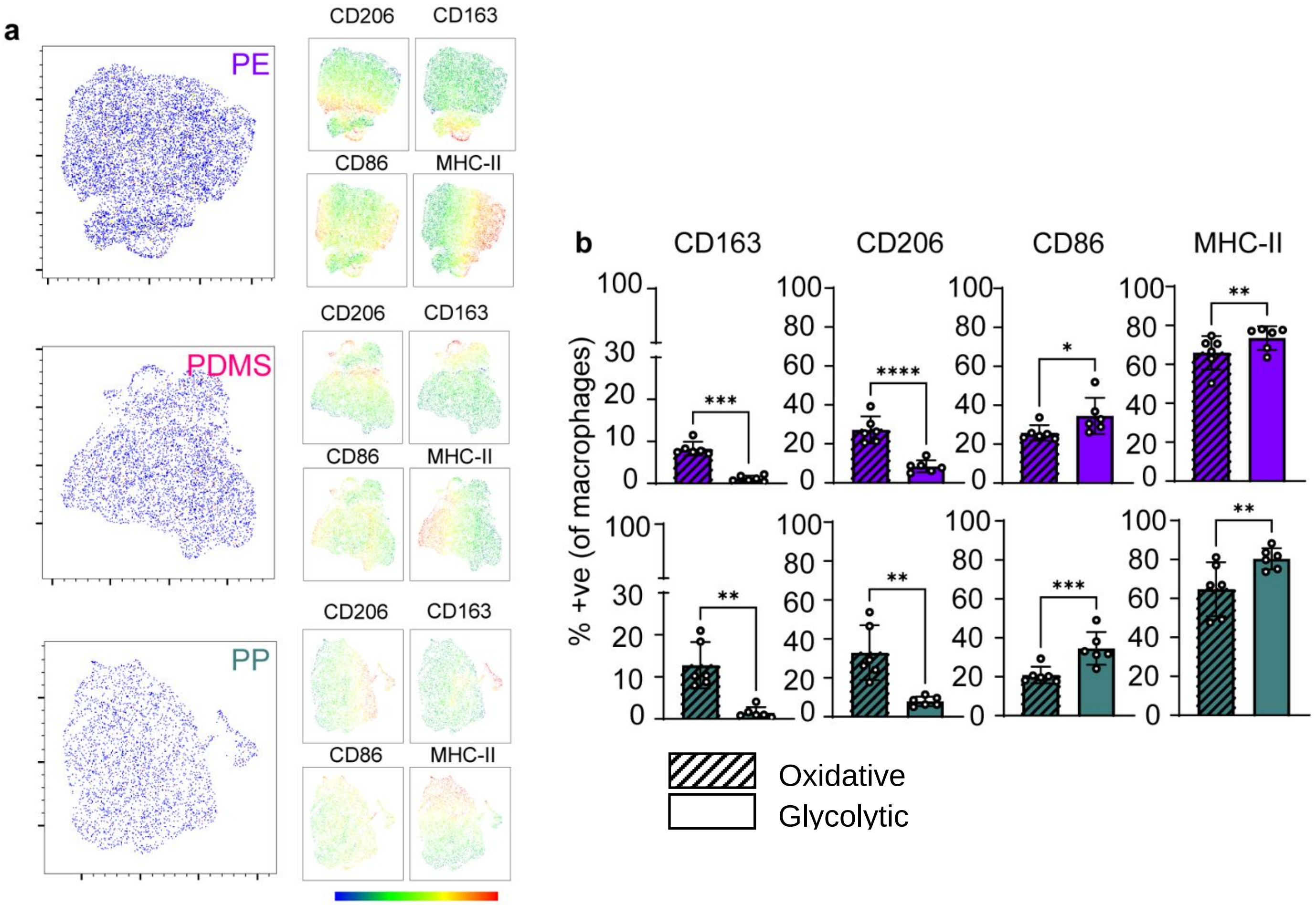


**Supplemental Figure 4: Expression of canonical macrophage phenotypic markers and their correlation to metabolic utilization 6-WPI**. **a.** Two-dimensional representation **(**UMAPs) of peri-implant macrophages canonical marker expression at 6-WPI. **b.** Frequency of positive expression in glycolytic and oxidative macrophages within the oligomycin A treated group for PE and PP-associated macrophages. Hashed bars represent oxidative cells, solid bars represent glycolytic cells. n = 6. Compared using a paired t-test. *$p < 0.05$, **$p < 0.01$, ***$p < 0.001$, ****$p < 0.0001$.

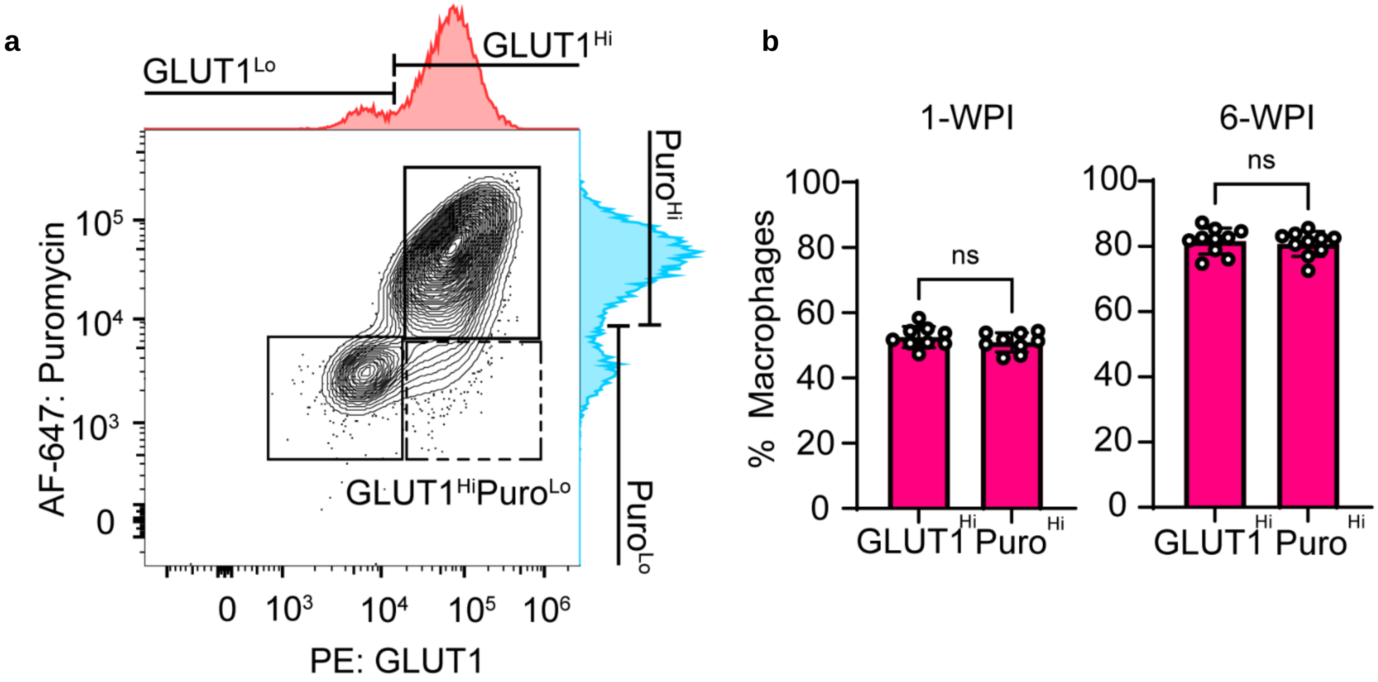


**Supplemental Figure 5: GLUT1$^{HI}$ cells are directly correlated with glycolytic utilization (Puromycin$^{Hi}$) in PDMS-associated macrophages at 1- and 6-WPI. a.** Representative co-expression of Puromycin and GLUT1 in an Oligomycin A treated sample indicate glycolytic and oxidative macrophage populations with consistently high and low GLUT1 expression (1) and a population highly expressing GLUT1 but with low puromycin intensity (2), which is not present in the control-treated samples. **b.** Frequency of GLUT1$^{Hi}$ macrophages and Puromycin$^{Hi}$ macrophages in matched vehicle and oligomycin A treated PDMS samples at 1- and 6-WPI. n = 9-10. Compared using a paired t-test. **p* < 0.05.

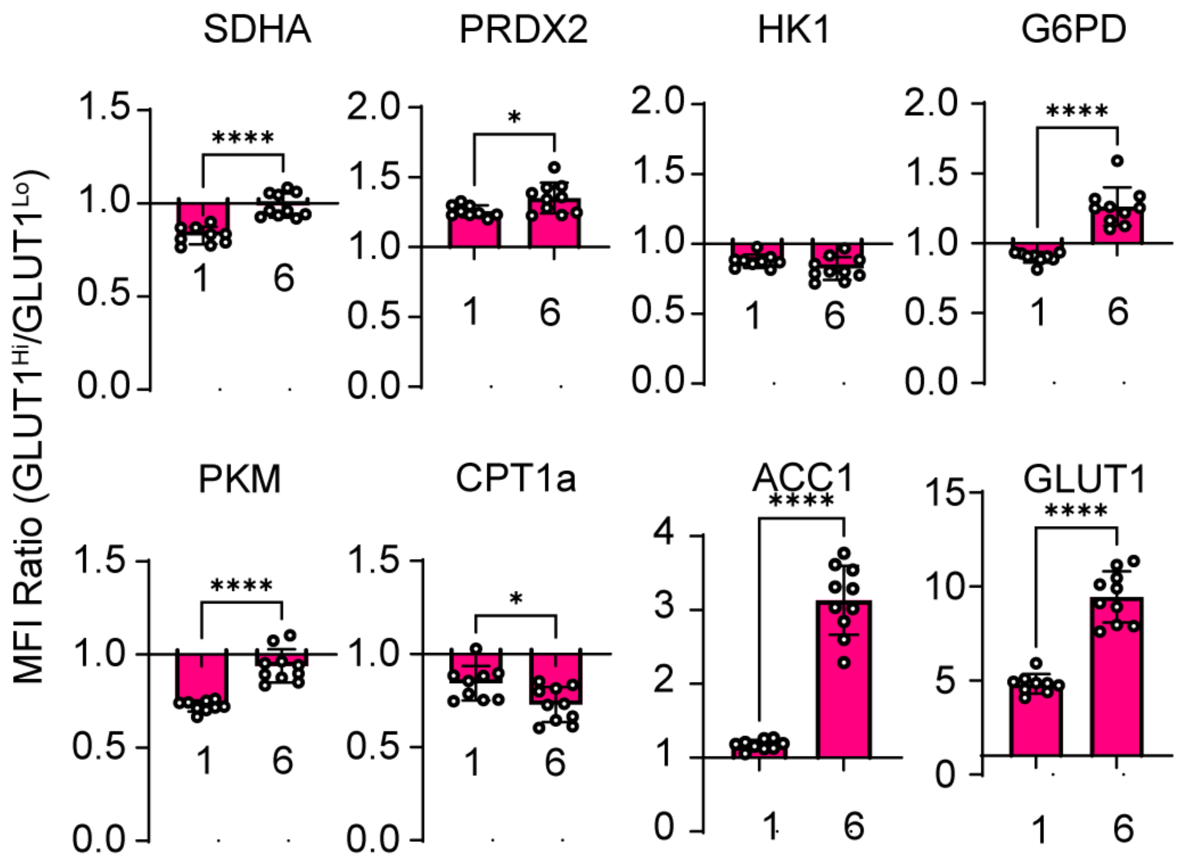


**Supplemental Figure 6: Ratio of metabolic enzyme expression in GLUT1^Hi^ to GLUT1^Lo^ cells**. n = 9-10. Compared using an unpaired t-test. *$p < 0.05$, **$p < 0.01$, ***$p < 0.001$, ****$p < 0.0001$.

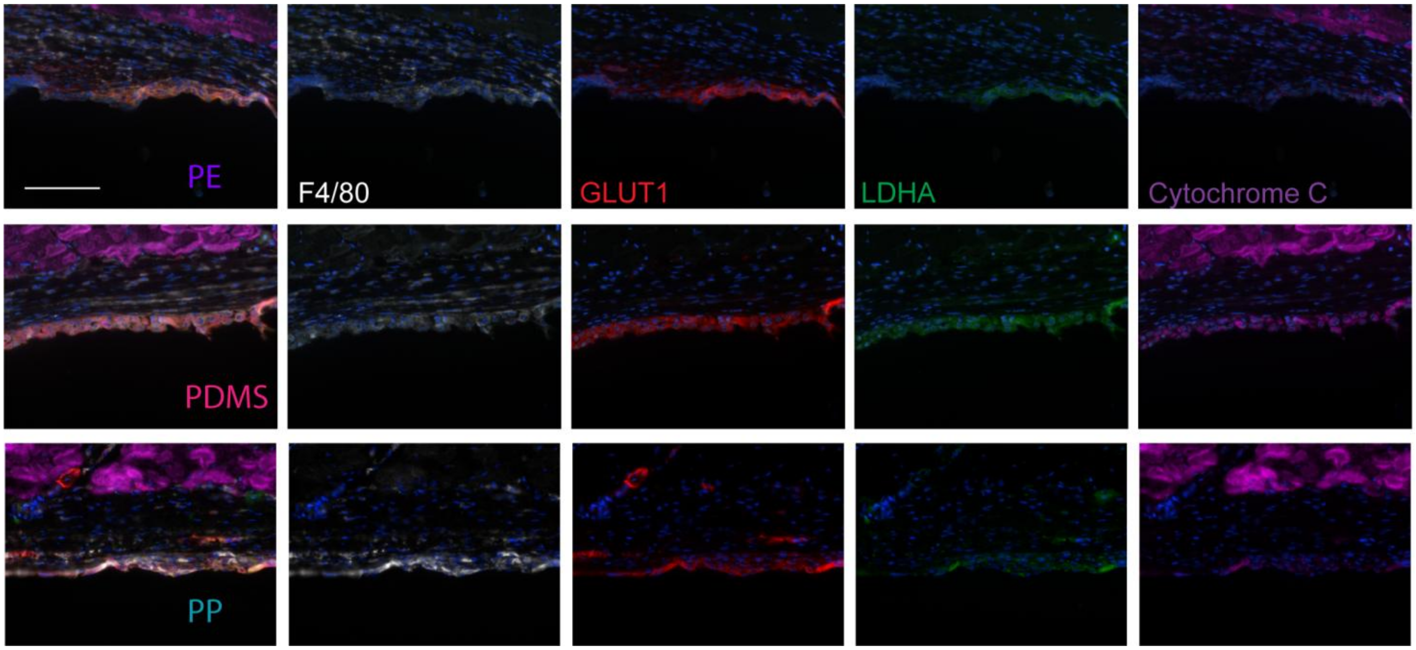


**Supplemental Figure 7: Representative spatial expression of metabolic markers in implant-associated tissues 6-WPI.** Composite images (left) and individual channels for PE, PDMS, and PP material image at 40X magnification. Scale bar = 100 µm.

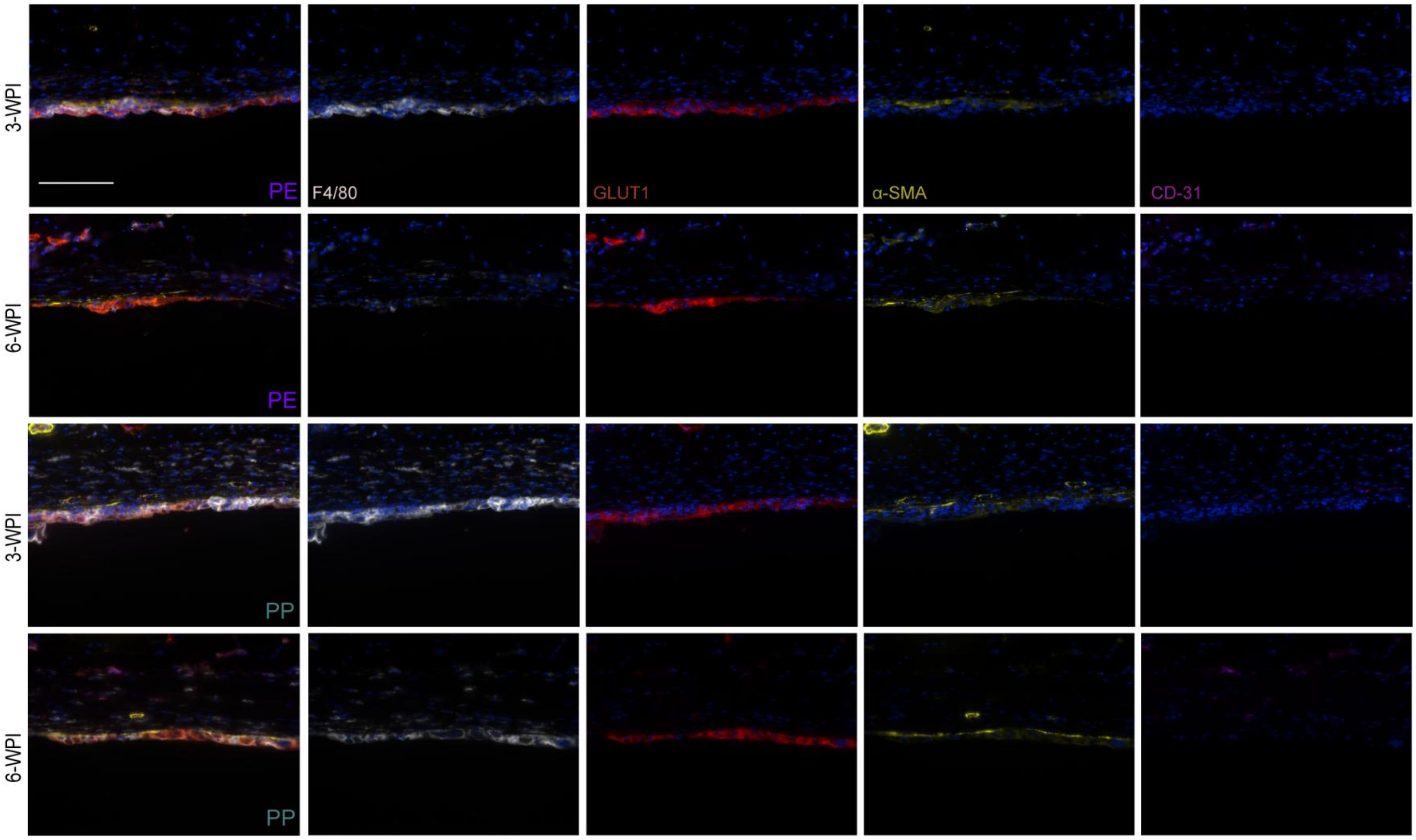


**Supplemental Figure 8: Representative OPAL™ Immunohistochemical staining of fibrotic markers in PE- and PP-associated tissues 3- and 6-WPI.** Presented as composite images and individual channels**.** Imaged at 40X magnification. Scale bar =100 µm.

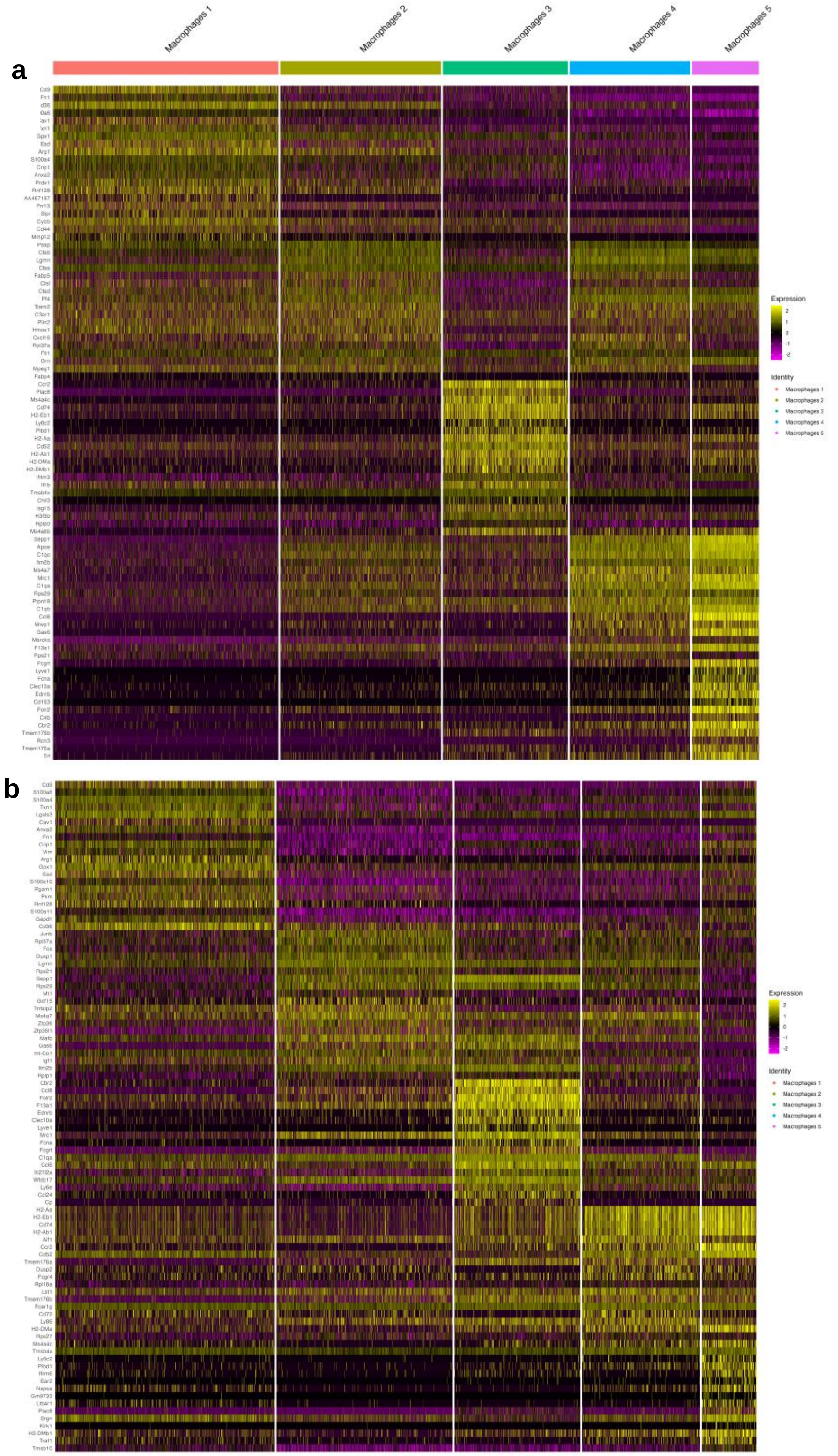


**Supplemental Figure 9: Heatmap of top 20 DEGs in scRNA-seq dataset for PDMS-associated macrophages** at 2-WPI (**a**) and 4-WPI (**b**). Relative expression normalized to the total number of transcripts in each cell.

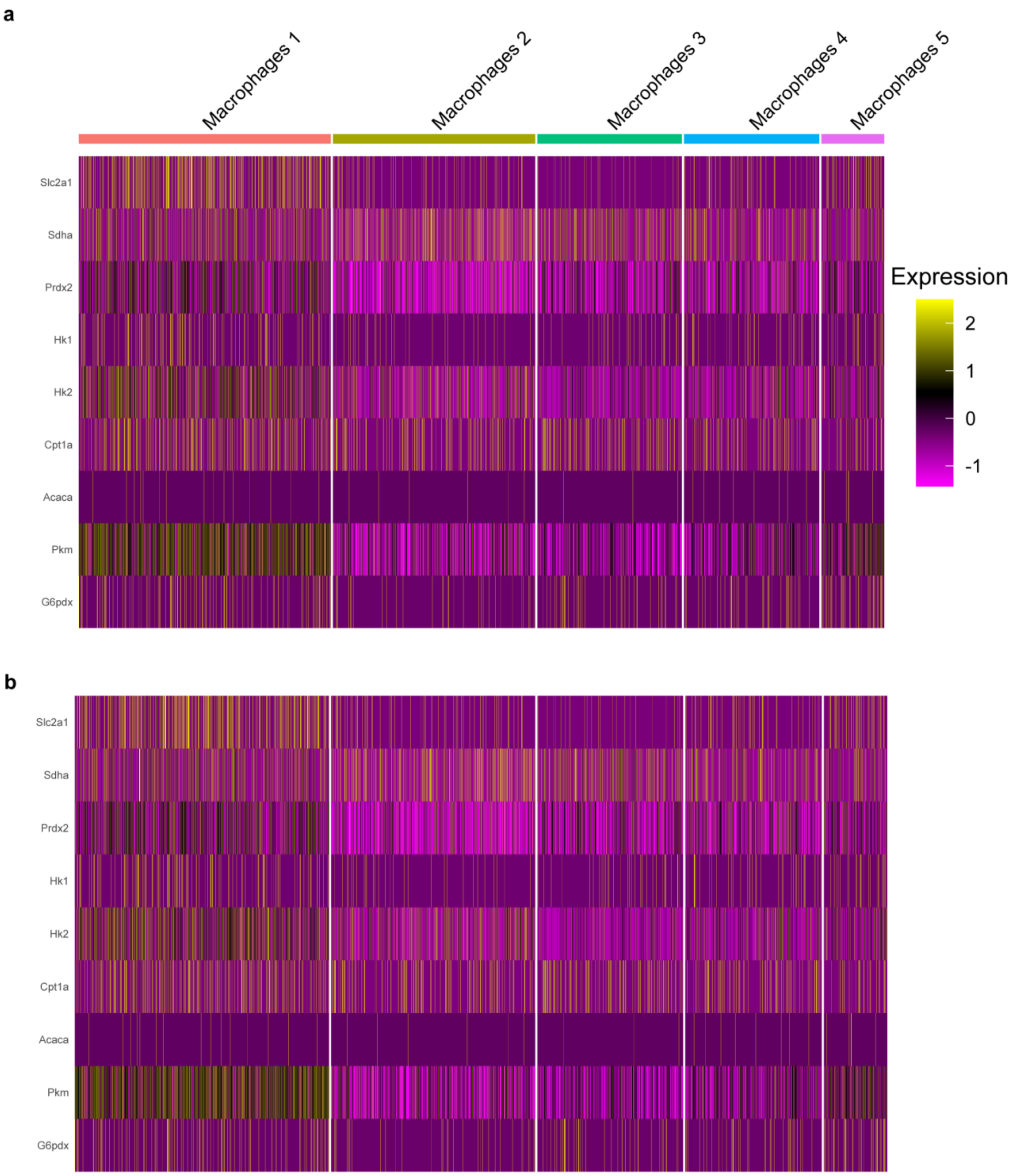


**Supplemental Figure 10: Heatmap of metabolic genes from scRNA-seq dataset for PDMS-associated macrophages** at 2-WPI (**a**) and 4-WPI (**b**). Relative expression normalized to the total number of transcripts in each cell

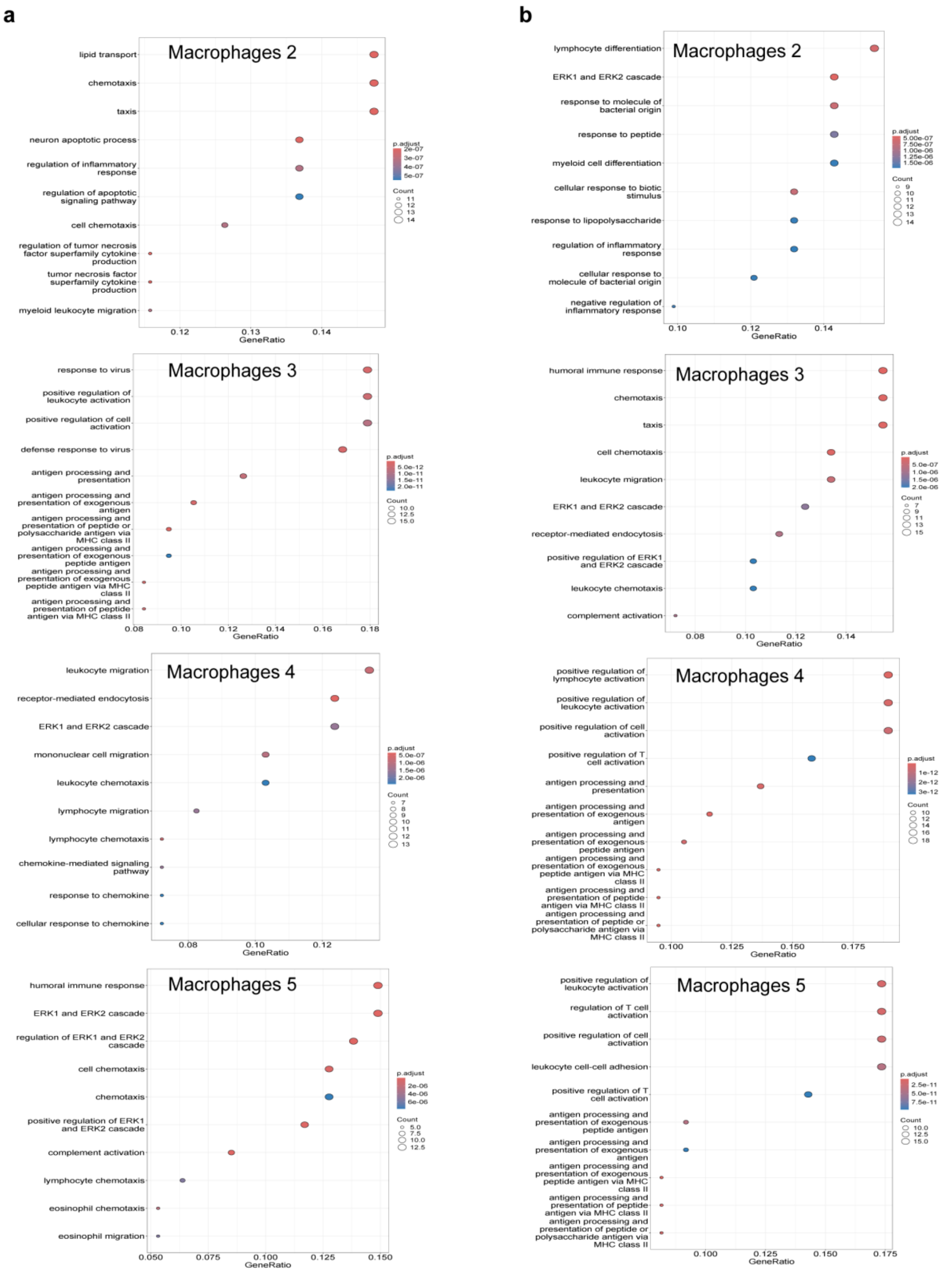


**Supplemental Figure 11: Dotplots representing gene ontology (GO) analyses in macrophage clusters 2-5 at 2- (a) and 4-WPI (b).** Top 10 pathways selected based on p-adjusted value and ranked based on gene ratio.

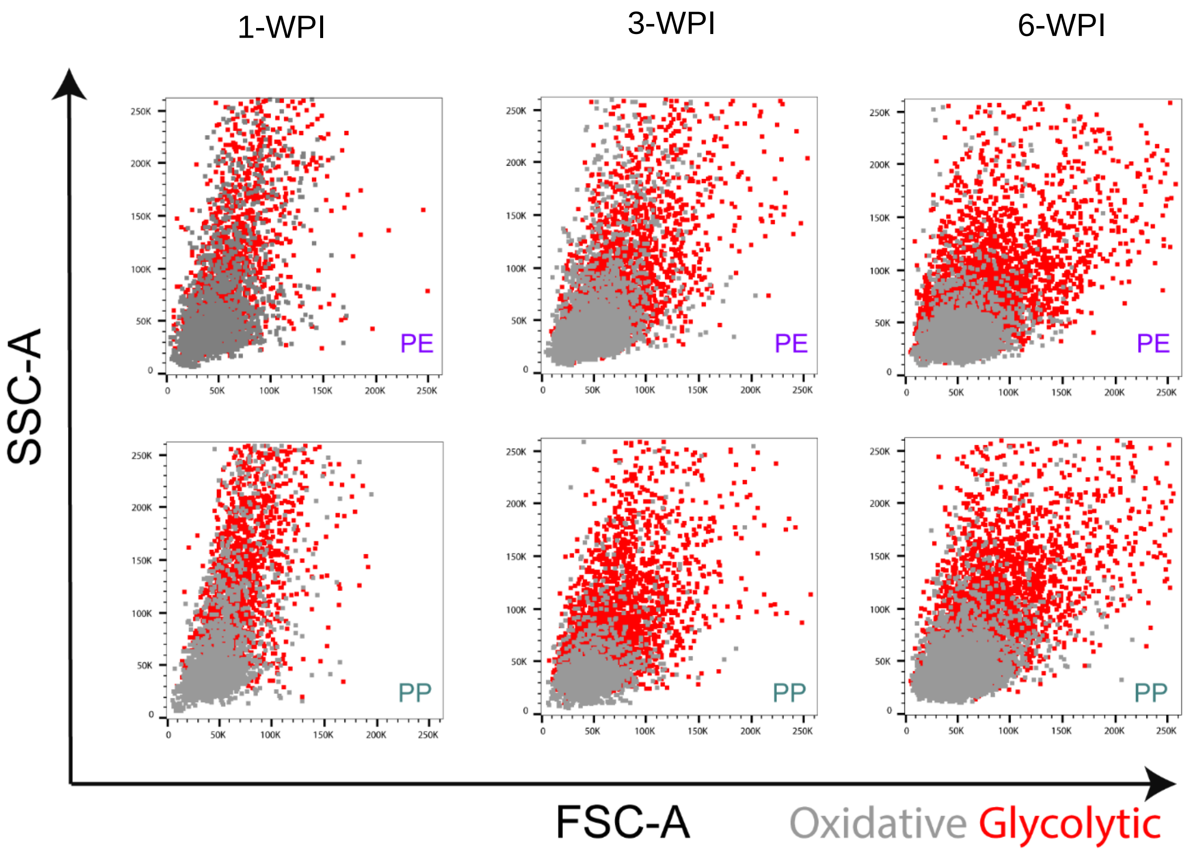


**Supplemental Figure 12: Representative scatter plots of glycolytic and oxidative macrophage size at 1-, 3- and 6-WPI in for PE and PP.**

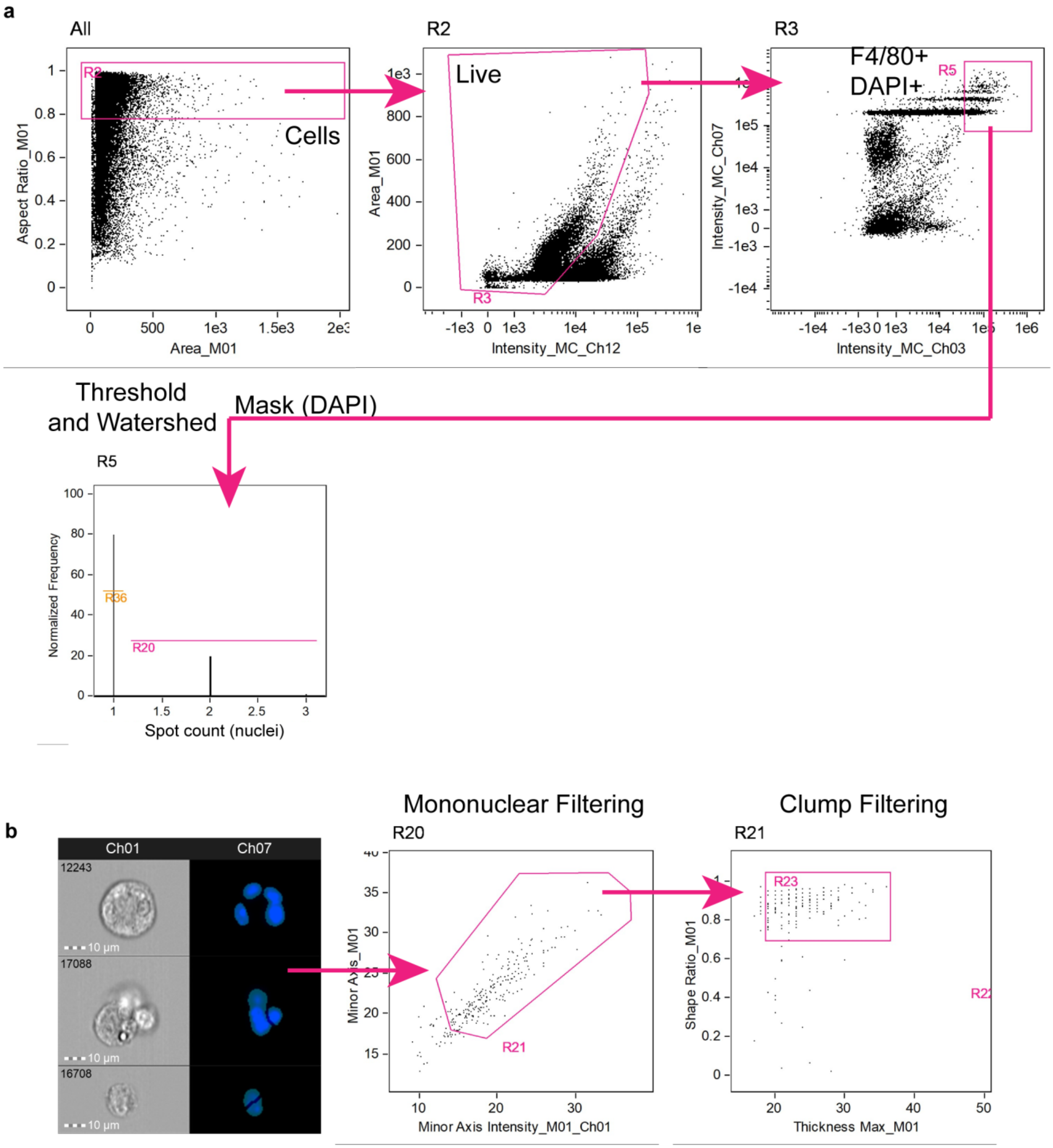


**Supplemental Figure 13: Gating scheme used for ImageStream® cytometry panel. a.** Gating scheme used to identify macrophages and multinucleated cells. **b.** Post-multinucleation gate processing to identify erroneously masked mononuclear cells and cell clumps.

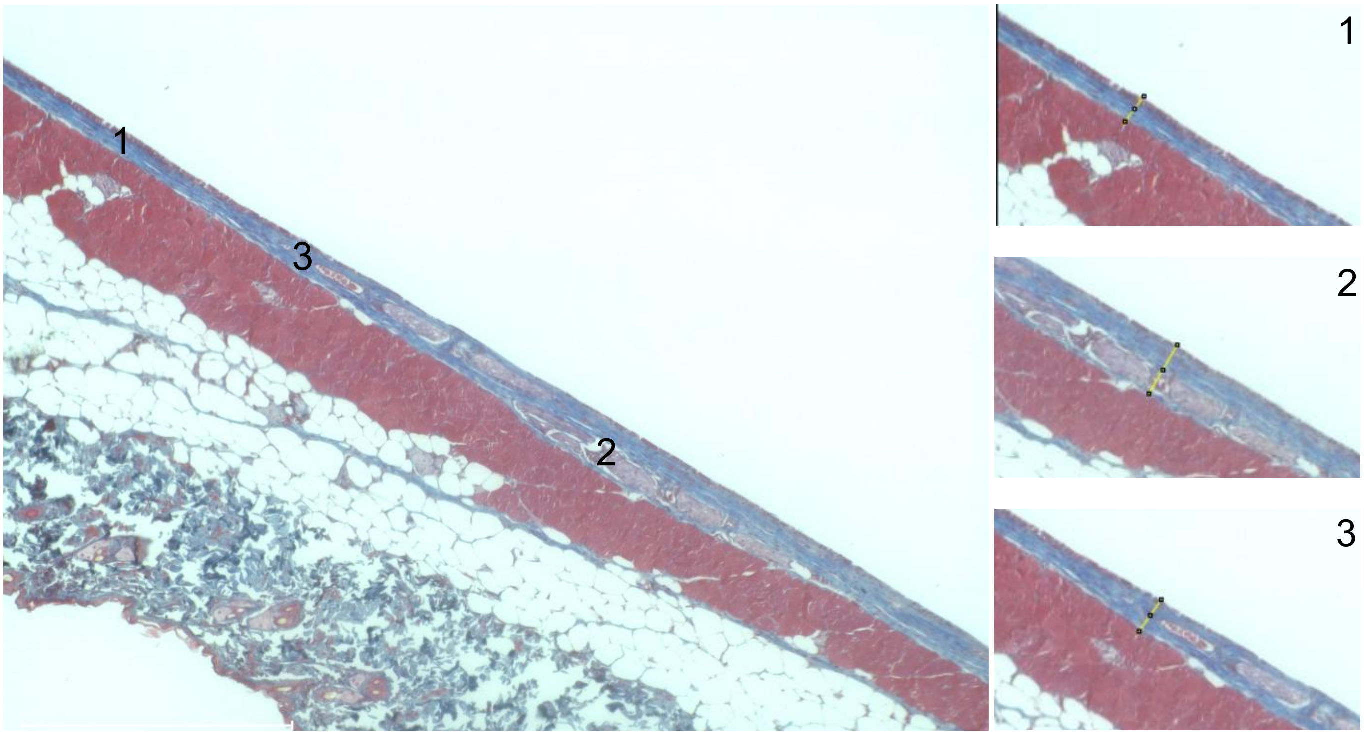


**Supplemental Figure 14: Representative imagery of the quantification method for Masson's trichrome stained slides.** Images were captured at 10X magnification and measured using ImageJ2 at the least fibrotic (1), most fibrotic (2) and most 'representative' (3) points. Muscle tissue (red), collagen (blue) and nuclei (brown/black).

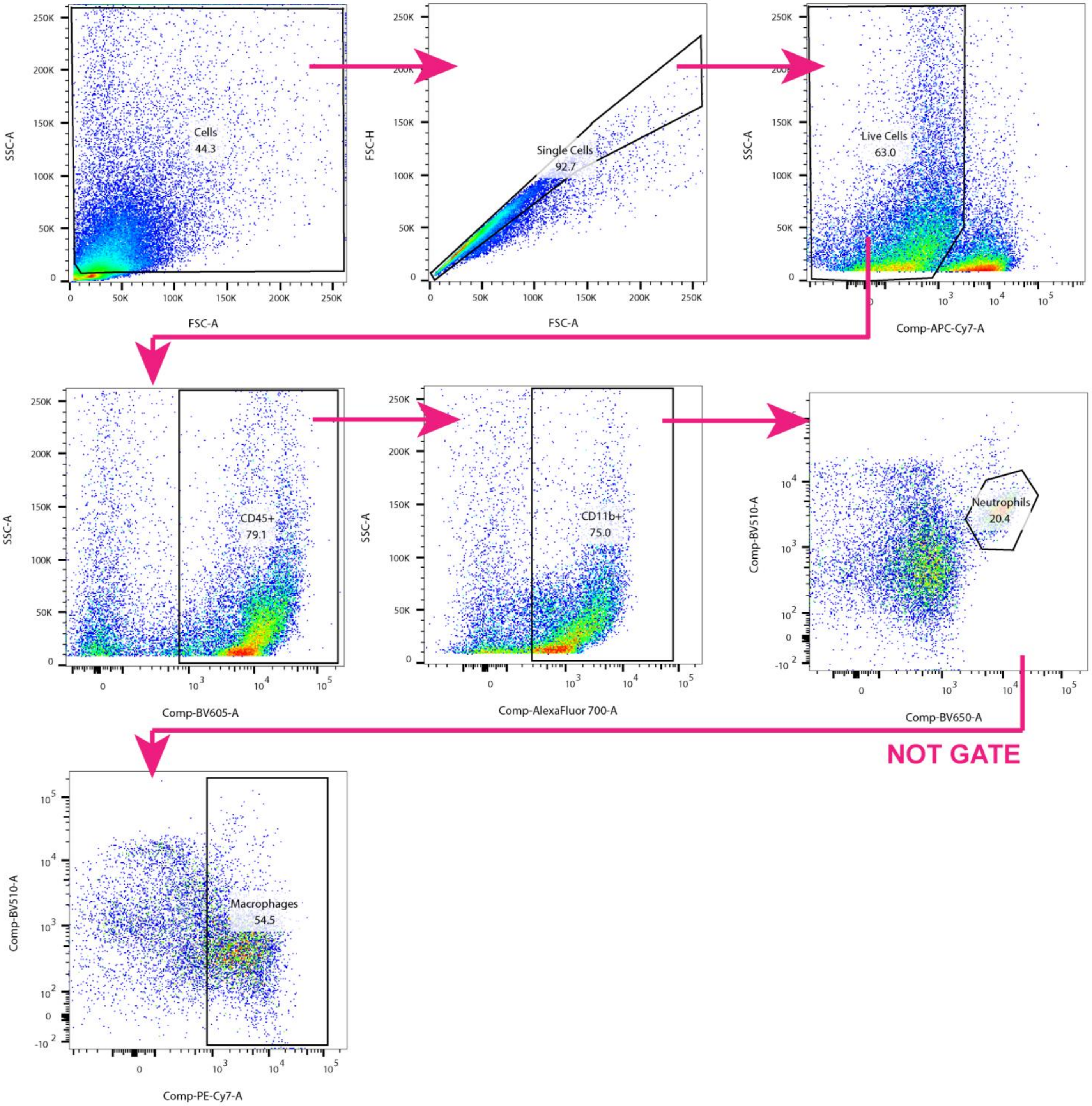


**Supplemental Figure 15: Gating scheme used for leukocyte frequency flow cytometry panel.**

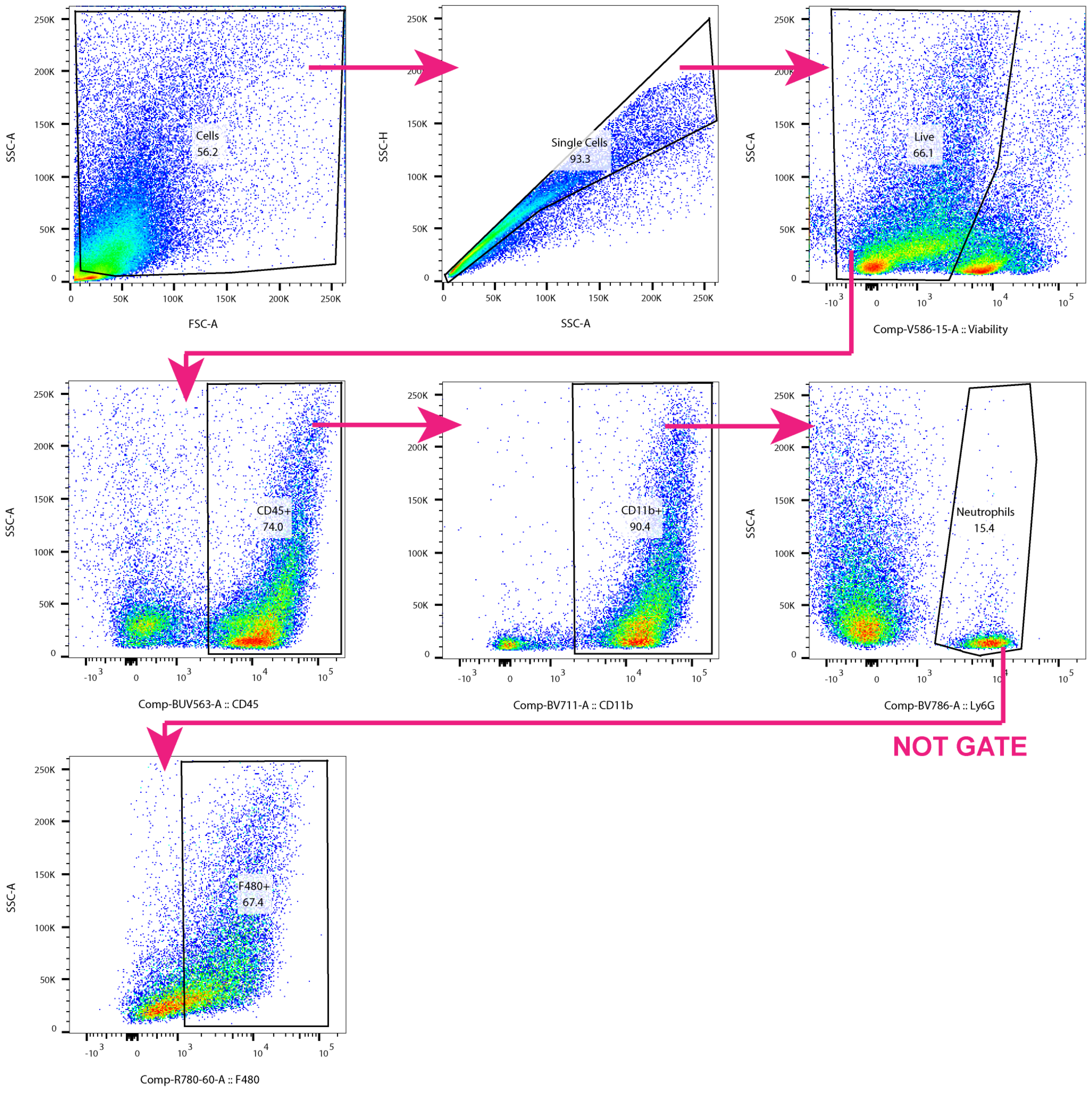


**Supplemental Figure 16: Gating scheme used metabolic enzyme expression flow cytometry panel (Met-flow).**

**Supplemental Table 1. Target genes and primer sequences for bulk gene expression analyses**

| Target Gene | Forward Sequence (5’ -> 3’) | Reverse Sequence (3’ -> 5’) | Source |
|---|---|---|---|
| *Canx* | ATGGAAGGGAAGTGGTTACTGT | GCTTTGTAGGTGACCTTTGGAG | Thermo Fisher Scientific |
| *Rer1* | GGCTGGACAAGTCTACCCC | GCGTAGGTCACAATGTACCAAC | Thermo Fisher Scientific |
| *Col1a1* | GCTCCTCTTAGGGGCCACT | CCACGTCTCACCATTGGGG | Thermo Fisher Scientific |
| *Col3a1* | CTGTAACATGGAAACTGGGGAAA | CCATAGCTGAACTGAAAACCACC | Thermo Fisher Scientific |
| *Fn1* | GCTCAGCAAATCGTGCAGC | CTAGGTAGGTCCGTTCCCACT | Thermo Fisher Scientific |
| *Il1β* | AGCTCCAAGAAAGGACGAACA | GCCCTGTAGGTGAGGTTGAT | Thermo Fisher Scientific |
| *Il6* | CCACAGTCCTTCAGAGAGATACA | CCTTCTGTGACTCCAGCTTAT | Thermo Fisher Scientific |
| *Tgfβ* | CGAAGCGGACTACTATGCTAAA | TCCCGAATGTCTGACGTATTG | Thermo Fisher Scientific |
| *Nos2* | GTTCTCAGCCCAACAATACAAGA | GTGGACGGGTCGATGTCAC | Thermo Fisher Scientific |
| *Adgre1* | TGACTCACCTTGTGGTCCTAA | CTTCCCAGAATCCAGTCTTTCC | Thermo Fisher Scientific |
| *Tnf* | GCCTCTTCTCATTCCTGCTT | TGGGAACTTCTCATCCCTTTG | Thermo Fisher Scientific |

**Supplemental Table 2.** Antibodies used for flow cytometry.

| Antibody Target | Clone | Fluor | Supplier | Product# | Dilutions |
|---|---|---|---|---|---|
| Anti-ACC1 | EPR23235-147 | Unconjugated | ABCAM | AB272704 | 1/1000 |
| Anti-CD11b | M1/70 | AF700 | BioLegend | 101222 | 1/500 |
| Anti-CD11b | M1/70 | BV711 | BioLegend | 101242 | 1/500 |
| Anti-CD45 | 30-F11 | BV605 | BioLegend | 103155 | 1/750 |
| Anti-CD45 | 30-F11 | BUV563 | BD Biosciences | 741274 | 1/500 |
| Anti-CD86 | GL-1 | BV421 | BioLegend | 105032 | 1/200 |
| Anti-CD163 | S15049I | KB520 | BioLegend | 155317 | 1/200 |
| Anti-CD206 | C068C2 | PerCP-Cy5.5 | BioLegend | 141715 | 1/200 |
| Anti-CPT1a | EPR21842-71-2F | Unconjugated | ABCAM | AB235841 | 1/200 |
| Anti-F4/80 | BM8 | PE | BioLegend | 123110 | 1/750 |
| Anti-F4/80 | BM8 | PE-Cy7 | BioLegend | 123114 | 1/750 |
| Anti-F4/80 | BM8 | APC-Fire™ | BioLegend | 1213152 | 1/200 |
| Anti-F4/80 | BM8 | BV605 | BioLegend | 123133 | 1/200 |
| Anti-G6PD | EPR20688 | Unconjugated | ABCAM | AB231828 | 1/1000 |
| Anti-GLUT1 | EPR3915 | PE | ABCAM | AB209449 | 1/200 |
| Anti-HK1 | EPR10134(B) | BUV661 | BD Biosciences | 570557 | 1/50 |
| Anti-Ly6C | HK1.4 | BV510 | BioLegend | 128033 | 1/500 |
| Anti-Ly6G | 1A8 | BV650 | BioLegend | 127641 | 1/500 |
| Anti-Ly6G | 1A8 | BV785 | BioLegend | 127645 | 1/500 |
| Anti-Ly6G | 1A8 | AF488 | BioLegend | 127626 | 1/500 |
| Anti-MHC-II | M5/114.15.2 | BV785 | BioLegend | 107645 | 1/500 |
| Anti-SDHA | EPR9043(B) | Unconjugated | ABCAM | AB240098 | 1/1000 |
| Anti-PKM | EPR10138(B) | Unconjugated | ABCAM | AB206129 | 1/200 |
| Anti-PRDX2 | EPR5154 | BUV615 | BD Biosciences | 570740 | 1/50 |
| Anti-Puromycin | 12D10 | AF647 | Sigma-Aldrich | Mabe343-af647 | 1/1000 |

**Supplemental Table 3.** Antibodies used for OPAL™ fluorescent histology.

| Antibody Target | Clone | Fluor | Supplier | Product# | Dilutions |
|---|---|---|---|---|---|
| Anti-α-SMA | 1A4 | Unconjugated | ABCAM | AB7817 | 1/500 |
| Anti-CD31 | D8V9E | Unconjugated | CellSignaling | 77699S | 1/200 |
| Anti-CD68 | SP251 | Unconjugated | ABCAM | AB192847 | 1/200 |
| Anti-Cytochrome C | EPR1326-80-5 | Unconjugated | ABCAM | AB76107 | 1/250 |
| Anti-F4/80 | D2S9R | Unconjugated | CellSignaling | 70076 | 1/250 |
| Anti-GLUT1 | EPR3915 | Unconjugated | ABCAM | AB115730 | 1/250 |
| Anti-LDHA | 2E2G6 | Unconjugated | Proteintech | 66287-1-Ig | 1/100 |
